# The human ribosomal RNA gene is composed of highly homogenized tandem clusters

**DOI:** 10.1101/2021.06.02.446762

**Authors:** Yutaro Hori, Akira Shimamoto, Takehiko Kobayashi

**Affiliations:** Institute for Quantitative Biosciences, the University of Tokyo, Tokyo, Japan; Faculty of Pharmaceutical Sciences, Sanyo-Onoda City University, Sanyo Onoda, Yamaguchi, Japan

**Keywords:** ribosomal RNA gene (rDNA), rDNA copy number, DNA methylation, senescence, genome instability, human, mutation rate, Oxford Nanopore sequencer, gene conversion, homogenization, progeroid syndrome

## Abstract

The structure of the human ribosomal RNA gene clustering region (rDNA) has traditionally been hard to analyze due to its highly repetitive nature. However, the recent development of long-read sequencing technology, such as Oxford Nanopore sequencing, has enabled us to approach the large-scale structure of the genome. Using this technology, we found that human cells have a quite regular rDNA structure. Although each human rDNA copy has some variations in its non-coding region, contiguous copies of rDNA are similar, suggesting that homogenization through gene conversion frequently occurs between copies. Analysis of rDNA methylation by Nanopore sequencing further showed that all of the non-coding regions are heavily methylated, whereas about half of the coding regions are clearly unmethylated. The ratio of unmethylated copies, which are speculated to be transcriptionally active, was lower in individuals with a higher rDNA copy number, suggesting that there is a mechanism that keeps the active copy number stable. Lastly, the rDNA in progeroid syndrome patient cells with reduced DNA repair activity had more unstable copies as compared with control normal cells, although the rate was much lower than previously reported using a Fiber FISH method. Collectively, our results alter the view of rDNA stability and transcription regulation in human cells, indicating the presence of mechanisms for both homogenization to ensure sequence quality and maintenance of active copies for cellular functions.

## INTRODUCTION

The ribosomal RNA gene (rDNA) is the most abundant repetitive gene in eukaryotic cells. In the budding yeast *Saccharomyces cerevisiae*, the structure and function of rDNA have been well studied, establishing rDNA as a unique region in the genome. Each unit (9.2 kb) of rDNA includes two coding regions, 35S precursor ribosomal RNA (rRNA) and 5S rRNA genes, and two non-coding intergenic spacer regions (IGSs) between the genes. The units tandemly repeat (~150 times) in chromosome XII (Petes 1979)(Kobyashi et al., 1998). A unique feature of the yeast rDNA is that it has a system to maintain the quality and quantity of the sequence in order to fulfill the huge demand of ribosomes in the cell (Gangloff et al. 1996) (for review see Kobayashi 2011). The rDNA tends to lose copies through recombination between them because of their repetitive nature and highly activated transcription. To maintain quantity, therefore, the rDNA amplifies more copies when the number is reduced (Kobayashi et al., 1998). As a result, the rDNA is continually undergoing contraction and expansion, and thus is one of the most unstable regions in the genome (Kobayashi 2014).

A DNA binding protein, Fob1 is a key player in the amplification reaction (Kobayashi 2003). It induces recombination for amplification by inhibiting replication at the replication fork barrier (RFB) (Supplemental Fig. S1). The inhibition induces a DNA double strand break at a relatively high frequency, and the repair process increases the number of copies by unequal sister chromatid recombination (Weitao et al. 2003) (Burkhalter and Sogo 2004) (Kobayashi et al. 2004). Recombination is also regulated by non-coding transcription (E-pro transcription) through cohesion dissociation (Kobayashi and Ganley 2005). In terms of quality control of rDNA, the sequences are always homogenized; that is, a copy with mutation is excluded by a Fob1-dependent recombination mechanism, such as gene conversion and contraction of the copies (Ganley and Kobayashi 2007). In fact, the rDNA sequences in the budding yeast are known to be relatively uniform even though about half of the copies are not transcribed (Ganley and Kobayashi 2011). Therefore, we speculate that active recombination in the rDNA maintains the integrity that ensures intact rRNA and the ribosome (Kobayashi 2014).

There is another face of such unstable rDNA – namely, it induces cellular senescence in budding yeast (Ganley and Kobayashi 2014). For example, in the *fob1* mutant, the rDNA is stable with less recombination and the mutant’s lifespan is extended by ~60% (Takeuchi et al. 2003; Defossez et al. 1999). In contrast, in the *sir2* mutant, in which E-pro transcription is enhanced and the rDNA copy number frequently changes, lifespan is shortened by ~50% (Saka et al. 2013; Kaeberlein et al. 1999). Because the rDNA is a large unstable region in the genome, its instability may affect the stability of the whole genome and thereby influence lifespan (i.e., the rDNA theory of aging) (Kobayashi 2008).

While we have good knowledge about yeast rDNA and its extra coding functions for aging, there is limited information on human rDNA. One reason is that the human rDNA unit (~43 kb) is much larger than the yeast rDNA unit (~9.2 kb) and it includes many small repetitive sequences in the non-coding region. Although the Human Genome Project declared its completion in 2003, it was difficult to assemble the rDNA into its actual composition using the relatively short “reads” that were obtained from the sequencing technology of those days. However, the recent development of DNA polymerase-independent long read sequencing technologies, such as the Oxford Nanopore or PacBio systems, has made it possible to assemble complete sequences of the unexplored regions (Miga et al. 2020).

The human rDNA is comprised of 100~500 copies in a cell (Parks et al. 2018; Agrawal and Ganley 2018). Each unit of rDNA consists of the 45S precursor RNA gene (45S rDNA), whose transcript is processed into mature 18S, 5.8S, and 28S RNAs, and the IGS, which is filled with small repetitive sequences such as microsatellites and transposons (Fig. 1A). In the IGS, there are two typical repeats: the R repeat and Butterfly/Long repeat. The R repeat (~680 bp, typically three copies) is located in the termination region of the 45S rRNA gene. It contains the Sal box that is associated with the transcription factor TTFI (Grummt et al. 1986). TTFI, similar to yeast Fob1, functions to inhibit the replication fork to avoid the collision of RNA and DNA polymerase (Akamatsu and Kobayashi 2015). The Butterfly/Long repeat (~4,500 bp, typically two copies), which is composed of a Long repeat, CT microsatellite and Butterfly repeat, is located at approximately the center of the IGS (Fig. 1A) (Gonzalez and Sylvester 2001; Agrawal and Ganley 2018).

**Figure 1.**
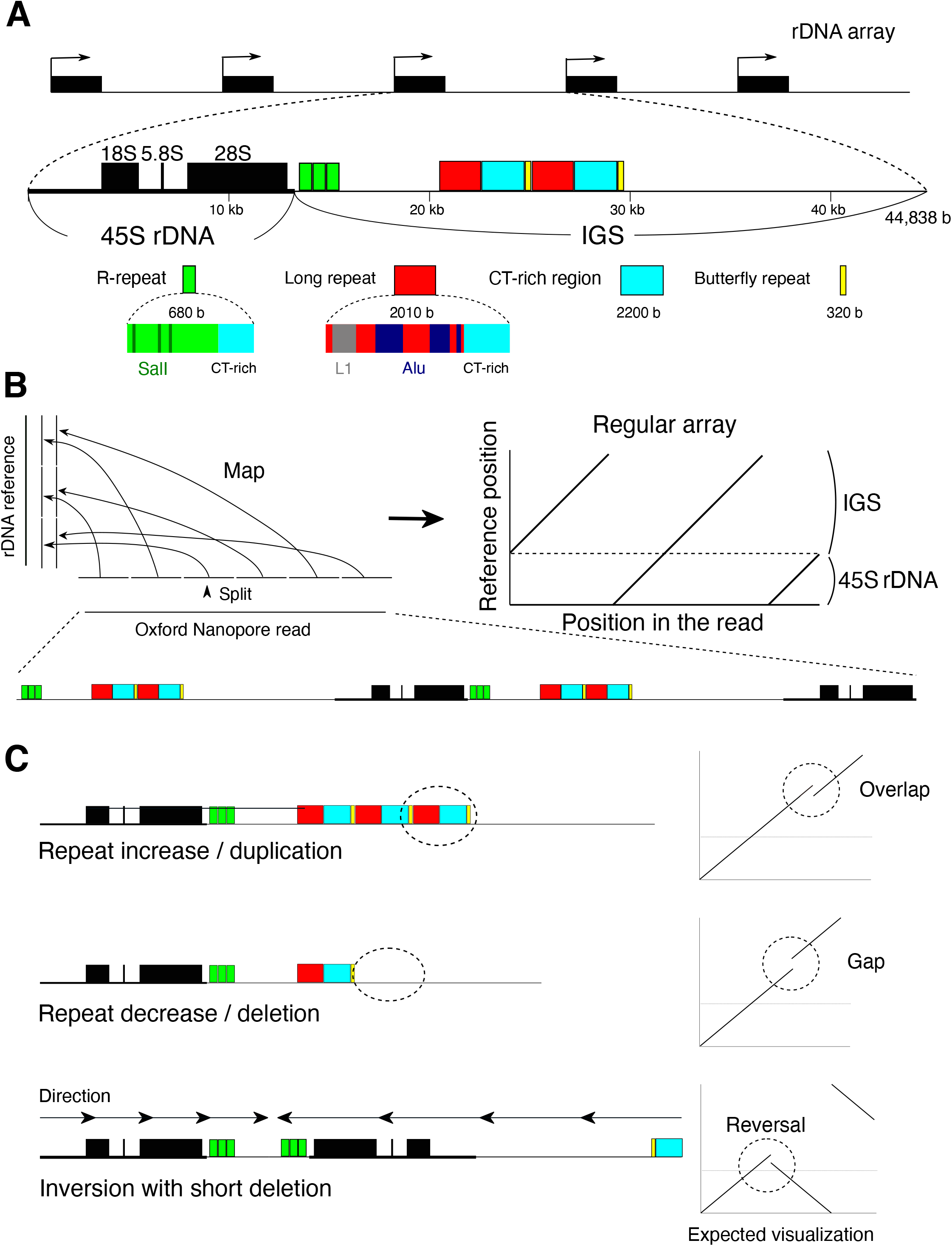
rDNA structure and strategy for visualizing rDNA. Structure of rDNA. rDNA is largely divided into the coding 45S rRNA gene (45S rDNA) and the non-coding intergenic spacer (IGS). The IGS has repetitive sequences, such as microsatellite and transposable elements. Here, typical R and Long/Butterfly repeats are shown. (B) rDNA visualization strategy for Nanopore reads containing rDNA. First, the read is split into 300-nt sections, and each split read is mapped to the rDNA reference sequence. The structure is then reconstructed based on the position in the read and the mapped position in the reference. (C) Typical mutations and how they look in the visualization strategy.

A previous study of rDNA composition in human cells by an *in situ* hybridization method (Fiber FISH) reported that many irregular units, such as palindromic inverted and incomplete units, account for ~35% of total copies in rDNA (Caburet et al. 2005). This high rate indicates that there is no effective recombination system to maintain rDNA homogeneity in human cells. In addition, the ratio of these non-canonical rDNA units was found to be increased in cells from progeroid syndrome patients, suggesting that human rDNA is also related to senescence. Because most progeroid syndromes, such as Werner syndrome and Bloom syndrome, are caused by mutations of the DNA repair machinery, it is plausible that the symptoms of these syndromes are caused by instability in the rDNA, which is thought to be one of the most unstable DNA regions in human, as in budding yeast (Carrero et al. 2016). Indeed, previous studies have suggested that rDNA copy number varies greatly in cells from Bloom syndrome patients, and palindromic structures have been observed in Werner syndrome cells (Schawalder et al. 2003; Killen et al. 2009). However, it is still unclear whether rDNA instability is an important factor in senescence in human. In terms of the relationship between rDNA and senescence in mammals, rDNA is also known to become methylated during the passage of life (Wang and Lemos 2019). The ratio of rDNA methylation works as an “clock” that tells the individual’s age.

To reveal the detailed structure and integrity in the human rDNA cluster at the DNA sequence level, here we developed a method to analyze rDNA-derived long reads obtained by an Oxford Nanopore sequencer. To our surprise, we found that the rDNA array is much more regular than we expected and the sequence similarity between adjacent copies is very high. These results suggest that recombination for homogenization takes place in the human rDNA as it does in the budding yeast.

## RESULTS

### Long sequence reads reveal variations among rDNA copies

To determine the human rDNA structure, we analyzed both publicly available Oxford Nanopore whole genome sequencing (WGS) data from the Human Pangenomics Project (HPGP, https://github.com/human-pangenomics/), and our in-house Cas9-enriched rDNA reads from Epstein-Barr Virus (EBV) transformed B cells (Shafin et al. 2020) and primary fibroblast cells (Supplemental Table S1). Taking advantage of the high copy number of rDNA per cell, we modified the Cas9-enrichment strategy and established a protocol to construct a library at lower cost (Gilpatrick et al. 2020). In short, to enrich the rDNA fragments, we designed four guide RNAs around the 9,500–9,900-nt region from the start site of the coding region (45S rDNA) of the reference sequence; all four sequences were strictly conserved in human and mouse. We designed gRNAs in the coding region because this region is thought to have fewer mutations and the relationship between two neighboring 45S rDNAs can be analyzed in a single read. Total DNA was dephosphorylated with CIP to avoid ligation to the sequencing adapter. The DNA was then digested by Cas9 ribonucleoprotein (RNP). Because only the Cas9-digested DNA fragments have a phosphorylated-5’ end, the sequencing adapters are specifically ligated to the fragments. In analyzing the HPGP genomic data, we removed reads shorter than 40,000 nt to eliminate those that were thought to come from rDNA-derived pseudo genes in non-rDNA genomic regions. For our in-house Cas9-enriched data, we analyzed the reads from DNA fragments in which both ends were digested with Cas9 RNP.

To determine the structure of rDNA, we developed a method that visualizes multiple copies of rDNA and the structural variation. In this method, the reads are split into 300 nt and mapped to an rDNA reference sequence (GenBank accession KY962518.1) using BWA MEM aligner software suited for long read mapping (Li 2013; Kim et al. 2018). The shorter split length increases not only resolution but also the effect of sequencing errors of the Nanopore sequencer and reduces mapping frequency. By testing several lengths of split reads, we found that a 300-nt split is long enough to accomplish high mapping frequency (Supplemental Fig. S2A). Each 300-nt split read (short line) is plotted based on its location in the original read and its mapped position in the reference sequence (see Materials & Methods; Fig. 1A, left panel). Therefore, when the reads are the same as the reference, a continuous straight line is generated (Fig. 1B, right panel). If there is a deletion, duplication or translocation, however, the line will be discontinuous (Fig. 1C). In selecting the reference sequence, we compared three different reference sequences of human rDNA (GenBank accession KY962518, U13369, and AL592188) by counting the number of gaps between successive split and mapped reads. If the distance between two neighboring mapped reads differed from the expected distance (300 nt) by more than 100 nt, we considered that the pair was gapped and we counted the number of gaps. We found that KY962518 had the least number of gaps for all samples and thus should be the most typical rDNA sequence as the reference (Supplemental Fig. S2B) (Agrawal and Ganley 2018; Kim et al. 2018).

Fig. 2A shows actual representative data of a Cas9-enriched read. The read (~40 kb) had one copy of rDNA with a gap in the Butterfly/Long repeat and a duplication in the R repeat region. Fig. 2B shows actual data from HPGP WGS sequencing. The length of this read was ~110 kb, corresponding to two and a half tandem copies of rDNA, that is, IGS–45S–IGS–45S–IGS. From the visualization, we could identify that all of the copies had the same duplicated regions in the Butterfly/Long repeat region. Therefore, this split-and-map visualizing method is robust and should work in the structural analysis of rDNA. A strong point of the long-read sequencer is that repeat structures of DNA can be analyzed; indeed, we were able to identify copies with extremely high repeat number variation in their IGS (Fig. 2C). In contrast, a weak point is that the fidelity of sequencing is not as high as that of short-read sequencers. But, as mentioned above, selecting an appropriate split length (300 nt) reduces this weakness and makes our structural analysis possible.

**Figure 2.**
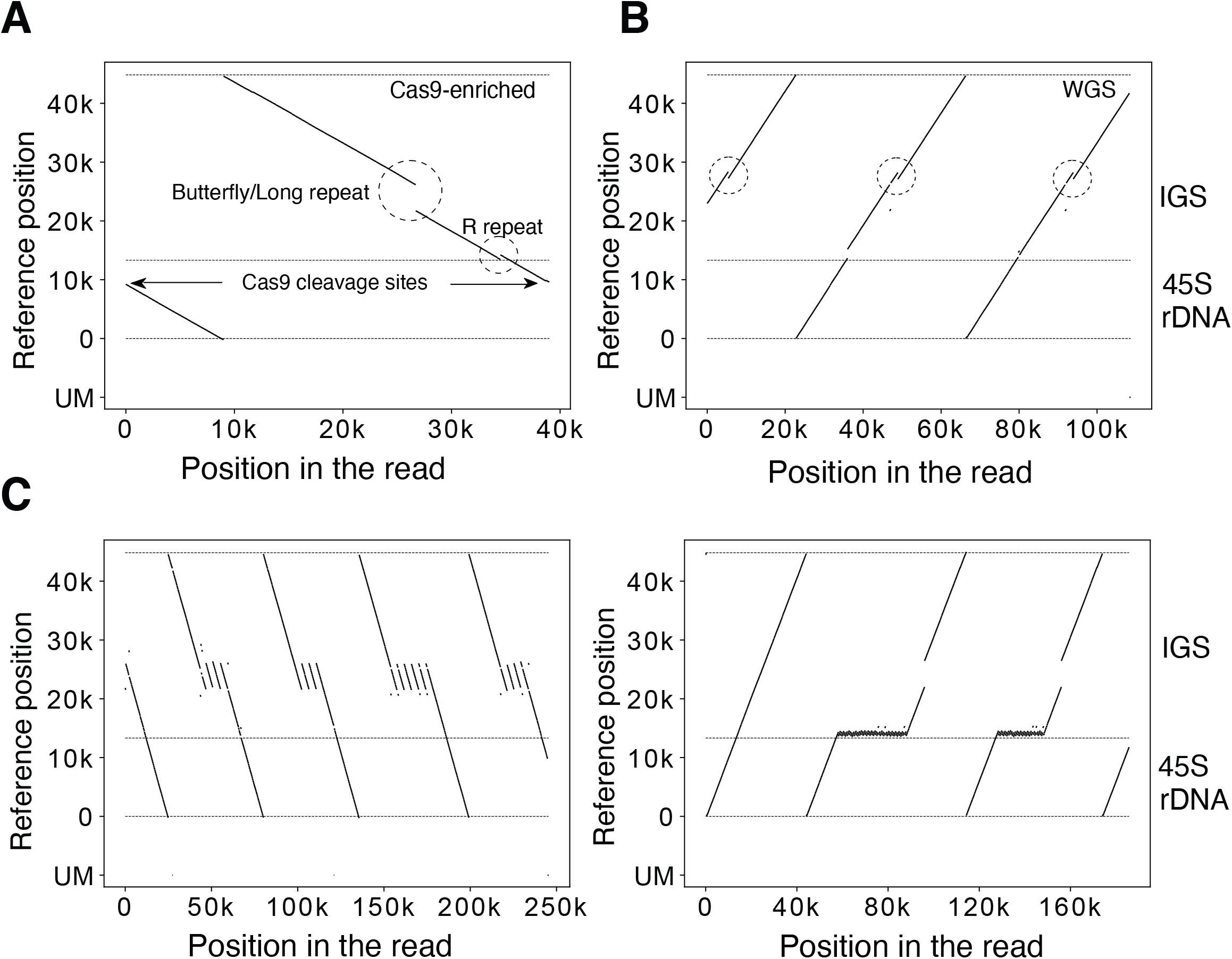
Visualization examples. (A)(B) Representative visualization pattern of a read obtained from Cas9-enrichment (A) and whole genome sequencings. The vertical axis is the mapped position in the reference; the horizontal axis is the position in the Oxford Nanopore read. The horizontal dashed lines indicate the end of the reference and the border between 45S rDNA and IGS. The unmapped split-reads are shown at the bottom (UM). The Sal box and the Butterfly/Long repeat region, which show variability among copies, are indicated (A). The same type of IGS variation is seen in all three rDNA copies (dashed circles) (B). (C) rDNA copies with an extremely long Butterfly/Long and R repeat. These sequences cannot be analyzed by short-read sequencers.

### R and Butterfly/Long repeats are highly variable between copies

By applying the split-and-map visualizing method to Nanopore data, we analyzed 39 samples (individuals) and identified variations in the Butterfly/Long and R repeat regions (Fig. 3A). By measurement of the Butterfly/Long repeat length of each read in each individual, we could classify the distribution into two types based on the proportion of copies that were more than 2,000 bases shorter than the reference (Fig. 3B; Supplemental Fig. 3A). These two types were also clearly differentiated by principal component analysis (PCA), confirming that our classification was not arbitrary (Supplemental Fig. S3B). In the shorter type, there were three discrete peaks and two of them were shorter than the reference. (e.g., HG03516). In the longer type (e.g., HG02080), the smaller peaks were not obvious in many cases, and more than 70% of copies were almost the same as the reference. Interestingly, all of the Japanese samples belonged to the shorter type (N=6: A0031, BSL2KA, PSCA0023, PSCA0047, PSCA0060, PSCA0517). Thus, there are differences among populations. In terms of the R repeat, the copy number varied from 0 to 4 (average 2–3 for most samples). As shown in Supplemental Fig. S4, the variation among samples was much larger for the R repeat than for the Butterfly/Long repeat and there were no clear differences among populations.

**Figure 3.**
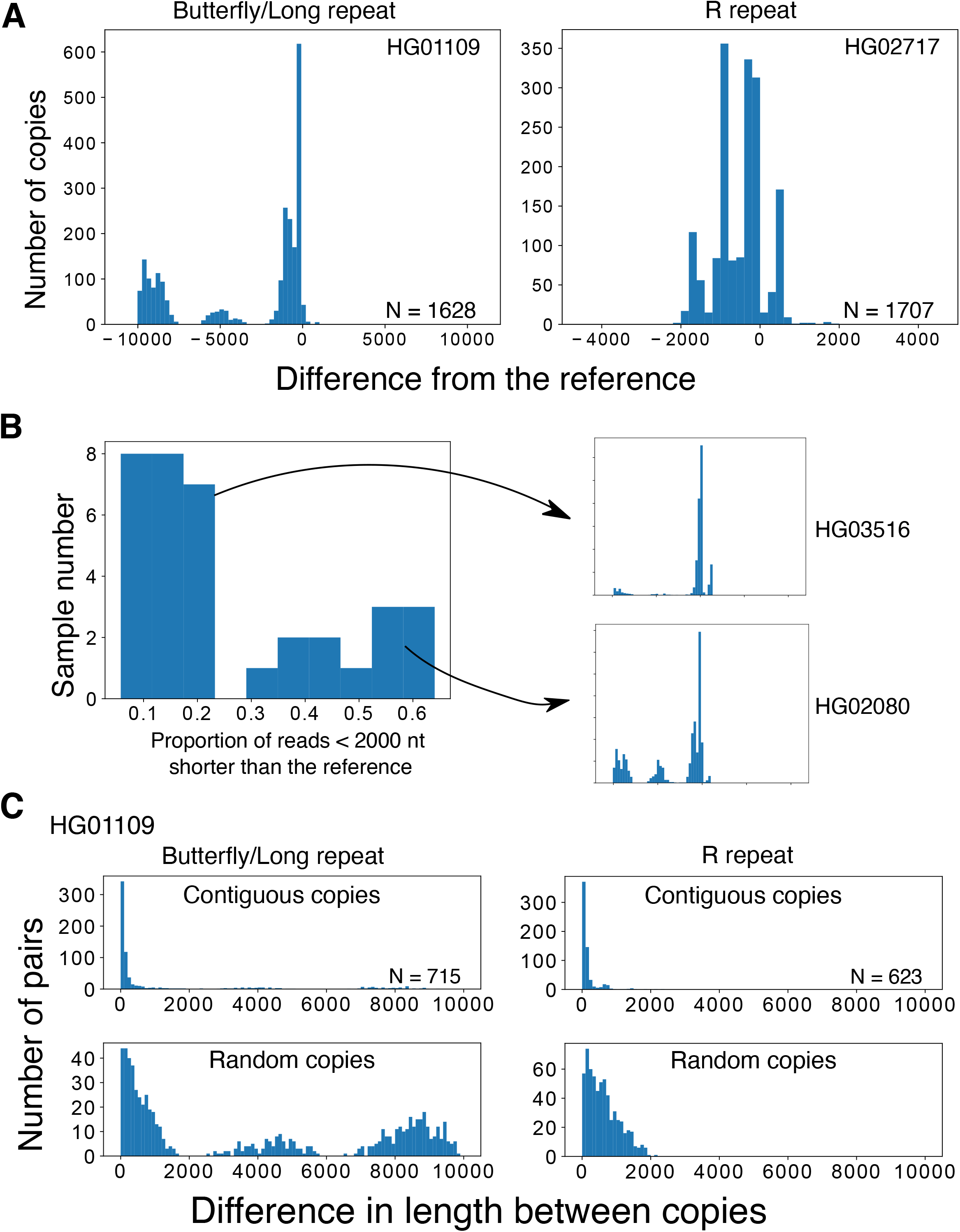
Variations in the IGS region. (A) Length distributions of the variable Butterfly/Long and R repeats of the rDNA IGS in WGS samples. Distributions are plotted based on the difference from the reference (0 indicates the reference length). In these samples, repeat number variations can clearly be seen in the discrete peaks. (B) Plot showing the proportion of reads with a Butterfly/Long repeat length shorter (<2,000 nt) than the reference for each sample. The samples can clearly be divided into two categories with each category having a similar pattern of size distribution among individuals (Supplemental Fig. 2). Typical distributions of long (HG03516) and short (HG02080) types are plotted on the right. (C) Differences in the length of contiguous reads calculated for the R and Butterfly/Long repeats (Upper panels). As a control, we calculated the differences in the length of the repeats of randomly picked copy pairs (Lower panels).

### Contiguous copies have similar variation patterns

By comparing copies within reads that contain more than one copy of rDNA, the similarity between contiguous copies can be calculated. Specifically, we analyzed the differences in lengths of the R and Butterfly/Long repeat regions (Fig. 3C, upper panel). As a control, we also simulated the case where two random copies are compared (Fig. 3C, lower panel; Supplemental Fig. S5). In all samples, the distributions of length difference between contiguous copies were clearly shorter than the randomized simulated control in both regions. These observations indicate that contiguous copies are more similar than non-contiguous ones. Therefore, this suggests that gene conversion, at least, occurs locally and homogenizes the sequences of these repeats.

### The rDNA is quite regular in human cells

Next, we analyzed larger-scale structural features of the rDNA. In a previous study, it was reported that human rDNA contains many non-canonical irregular copies, such as palindrome structures (Caburet et al. 2005). Such irregular structures were suggested by a Fiber FISH (DNA combing) assay, in which long rDNA spread on a slide glass is hybridized with two fluorescent probes for different sites in the rDNA and fluorescence signals are detected by microscopy. As a result, uncanonical copies are identified by the irregularity of the aligned dotted signals. The study indicated that irregular rDNA copies account for ~35% of total rDNA copies in a cell. Furthermore, this ratio was much increased in cells from Werner syndrome patients (~50%).

Using long sequence reads, we expected to obtain a more accurate description of the larger-scale structure of rDNA. First, we screened the reads in which more than 10% of the split reads were mapped in the opposite direction to the majority of the split reads and labeled them as inverted reads. Then, using the split-and-map visualizing data, we measured the distances between the splits, and compared them to the distance that was calculated from the reference sequence: if the repeat has an irregularly aligned or unusual structure, the distance between adjacent split reads will be larger and we can detect the difference. Unexpectedly, we found that reads with such large gaps are rare; that is, most of the rDNA copies were beautifully tandemly aligned on the chromosomes (Supplemental Table S1). In fact, the rate of non-canonical copies excluding palindromic reads in the healthy samples was less than 1% (0% ~ 0.7%). Furthermore, in ~30% (12/39) of the individual samples, no non-canonical copies were detected. The most common structural mutation was duplication from the R repeat to the Long repeat region (Fig. 3A left; Supplemental Table S1). We consider this type of mutation to be a common variation because it is limited to the IGS region, where it is likely to have little impact on rRNA transcription. Another typical structural mutation was deletion (Fig. 4A right).

**Figure 4.**
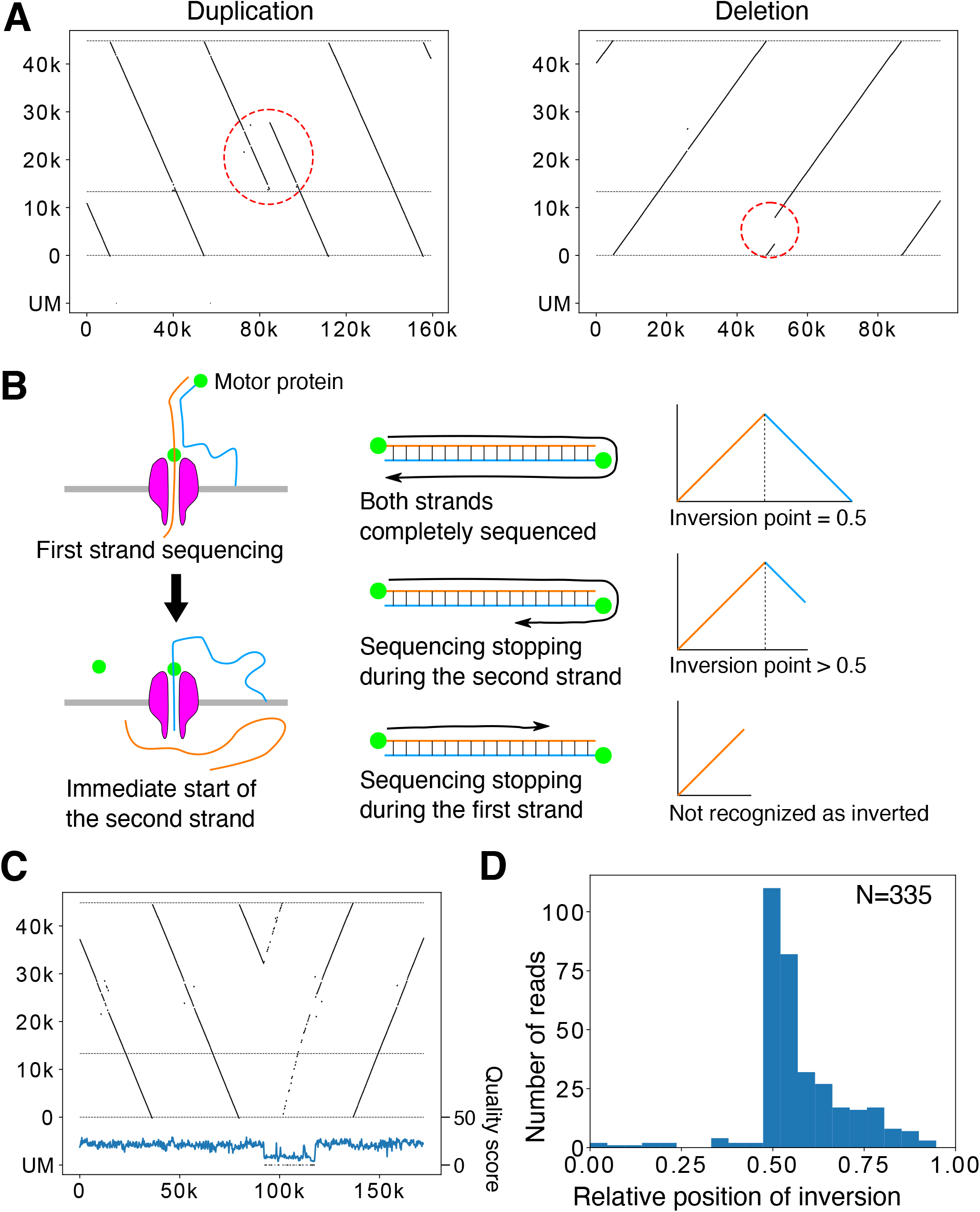
Large-scale structural variation in the rDNA array. (A) Representative reads with large-scale variation in the rDNA array. (Left panel) The R to Butterfly/Long repeat region is doubled. (Right panel) A large portion of rDNA in the 45S rDNA is deleted. (B) The mechanism of Nanopore template switching that causes artifactual “fake” palindromic reads. (Left panel) After the completion of first strand sequencing, second (complementary) strand sequencing sometimes occurs. (Right panel) When sequencing terminates randomly after strand switching, the resulting distribution of the inversion points in the reads should be seen in only the latter half of the reads. (C) A representative palindromic read. Note that the structure and the end points of the read are remarkably similar before and after the inversion point, although the read is relatively long. The Phred quality scores plotted below were smoothed by binning and averaging. A sudden drop in quality score is observed just after the inversion point. (D) Plot of the relative position of inversion in each read for HPGP samples. The distribution is peaked at the center and heavily skewed to the latter half of reads, suggesting that most of the palindromic reads are artifacts.

It should be noted that we found many palindromic reads, but they are thought to be artifacts for the following reason. The Oxford Nanopore sequencer reads single-stranded DNA by separating double-stranded DNA at the pore (Fig. 4B). If the separation does not occur properly at the end of the first strand, sequencing of the second complementary strand may follow immediately (de Lannoy et al. 2018). Therefore, the resulting sequence read will look like a palindrome. Such“fake” palindromic reads should have their inversion site at the center of the read in cases where they were sequenced completely. A sequencing reaction may stop at any point in a read for various reasons. If it stops after the inversion, the inversion site should be in the latter half of the sequenced read. Taken together, if a palindromic read is an artifact, the inversion site will be at the center or in the latter half of the read. We therefore investigated the relative position of the inversion site in each palindromic read for HPGP samples. As shown in Fig. 4D, most of the inversion points appeared after the center, which strongly indicates that most of the palindromic reads are the result of the aforementioned artifacts. In addition, many such pseudo-palindromic reads showed a sudden drop in sequencing quality score around the inversion site. Some of the reads with an inversion site in the former half also showed a sudden drop in sequencing quality score, possibly meaning that they also are not real palindromes. Nevertheless, assuming that the reads with their inversion site in the first half are true palindromes, the estimated frequency of palindromic inversion is ~1 in 2000 copies. In summary, the human rDNA is a very regular array (>99.3%) and aberrant structures such as palindromes are not common.

### 45S rDNA is “all-or-none-methylated”

Oxford Nanopore sequencing can also detect CpG methylation without any prior treatments. We therefore investigated the rate of CpG methylation in the human rDNA. The methylation status of each read was calculated by binning (bin size 200 nt). We used two methods for calculating the methylation frequency of 45S rDNA in each bin: first, we calculated the expected value of the proportion of methylated bases by simply taking the mean of the posterior probability of each CpG being methylated that was output by Oxford Nanopore Guppy basecaller (Supplemental Fig. S6 “average”); and second, we estimated the proportion of bases likely to be methylated by setting a threshold on the posterior probability (Supplemental Fig. S6, “threshold”). We obtained similar results by both methods; that is, the distinction between the methylated (>0.1) and less methylated (~0.0) copies was clear in most samples except A0031 and HG03098.

We present an example of the visualization of rDNA methylation level using this method in Fig. 5A. In this read, the right two 45S rDNA copies were less methylated and the left one was highly methylated. We summarized the methylation status of 45S rDNA in another sample (HG00733) (Fig. 5B). Clearly, there were methylated and less methylated copies of 45S rDNA. Notably, the less methylated 45S rDNA copies were almost methylation-free (“unmethylated”); therefore, we may say that 45S rDNAs can be classified in two states, summarized as “all-or-none-methylated” (Fig. 5B). Furthermore, the unmethylated copies are thought to be transcribed (Kass et al. 1997).

**Figure 5.**
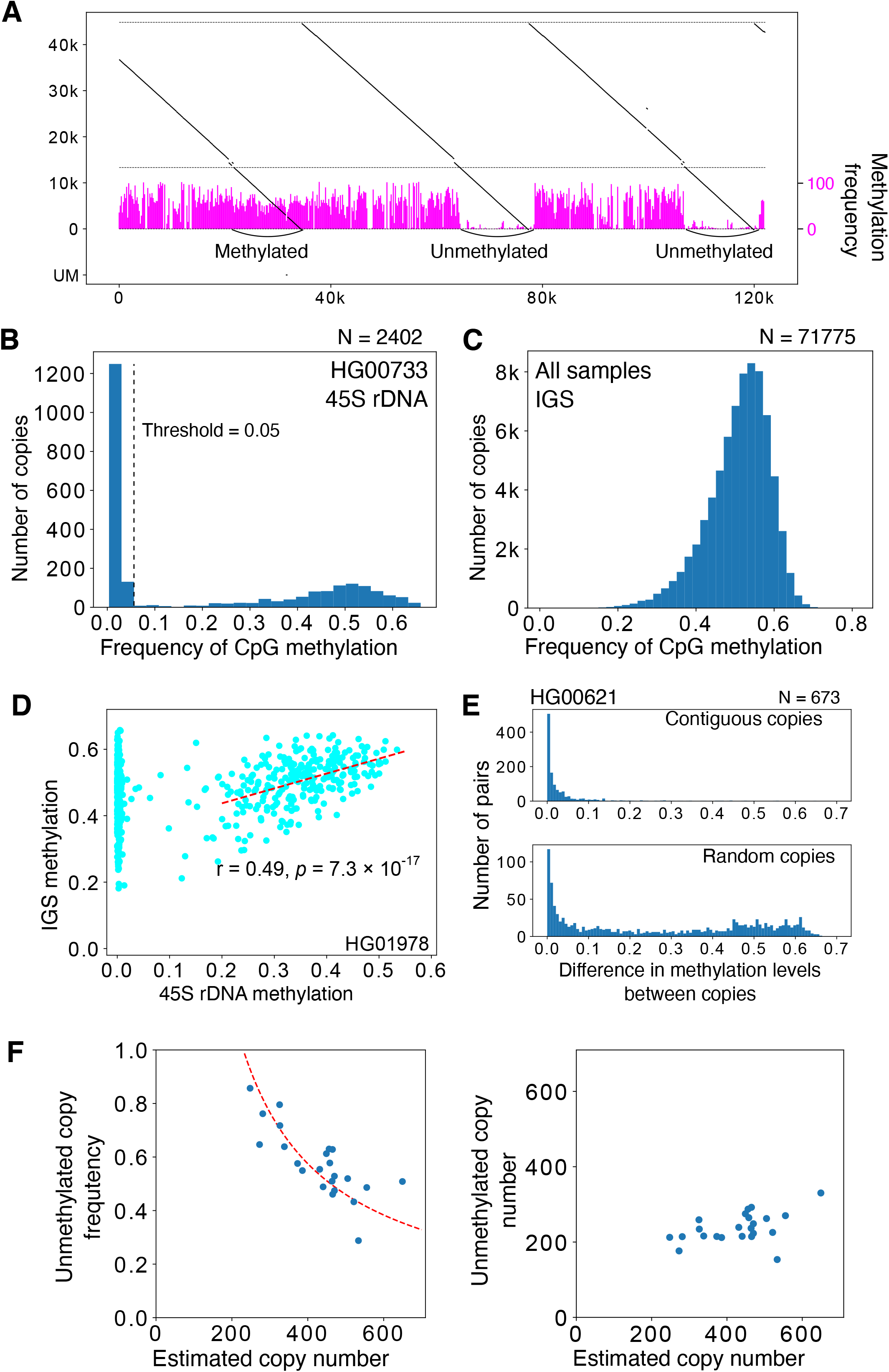
Methylation analysis of rDNA. (A) Representative visualization of CpG methylation. Reads are split into 200-nt bins and the expected frequency of CpG methylation is calculated for each bin using posterior probability output by Guppy basecaller. The methylation frequency of each bin is shown as a vertical blue bar. In the read shown, both the methylated and less methylated 45S rDNA are included. Note that the three IGS methylation patterns are similar despite the difference in the 45S rDNAs. (B) Average proportion of methylated CpG for each 45S rDNA (calculated by taking the mean of posterior probabilities), and distribution in the HG00733 sample. Dashed line indicates the border between the less methylated copies and methylated copies. (C) Proportion of CpG methylation in the IGS in all samples. Most of the IGS copies are heavily methylated. (D) Methylation level of 45S rDNA and its flanking IGS. In many samples, there is a clear correlation for copies with highly methylated 45S rDNA (dashed line). (E) Differences in the methylation levels of contiguous 45S rDNAs and randomized controls. The methylation pattern is similar between adjacent copies of contiguous 45S rDNA, as in the case of repeat number variation in the IGS (Fig. 3C). (F) Relationship between the estimated rDNA copy number and the proportion of unmethylated copies (left) or the estimated number of unmethylated copies (right). Dashed line in the left panel is a theoretical line based on the assumption that the number of active rDNA copy is constant at 230.

### Methylation in contiguous 45S rDNA and the IGS is correlated

In contrast to coding 45S rDNA copies, the non-coding IGS seems always to be methylated based on our visualization (Fig. 5A). Therefore, we investigated methylation status in the IGSs quantitatively. Because the Butterfly/Long repeat contains microsatellites with few CpG pairs and its length is variable, we excluded this region from the analysis. As a result, we found that nearly all of the IGSs are heavily methylated (average 51%) (Fig. 5C; Supplemental Fig. S7). Furthermore, we evaluated the methylation rate of 45S rDNA and its contiguous IGS and found that the rate of methylated bases in the IGS was correlated with the methylation level of contiguous 45S rDNAs in many individuals when the calculation was limited to strongly methylated 45S rDNA (>0.3), although the strength of correlation differed among samples (Fig. 5D; Supplemental Table S2). Overall, these observations suggest that heavily methylated 45S rDNA and the contiguous IGS together form heterochromatin.

### Contiguous 45S rDNAs have a similar CpG methylation pattern

Using information in large reads containing several rDNA copies, we analyzed the relationship of methylation status among contiguous 45S rDNAs (Fig. 5E; Supplemental Fig. S8). Interestingly, we found that contiguous 45S rDNA copies have a similar CpG methylation pattern as compared with noncontiguous random copies. In other words, unmethylated 45S rDNAs form clusters. This suggests that heterochromatin structure is present in the large rDNA region and the transcription of rDNA is inhibited in this region.

We also examined how often methylation status changes in each chromosome. A previous study suggested that the transcriptional state of rDNA is determined at the chromosome level (Roussel et al. 1996). We found that a contiguous pair with different methylation status occurs at the frequency about 1 in 20 pairs in many individuals (Supplemental Table S3). Therefore, chromosomes that have more than 20 copies of rDNA should have, on average, more than one change in methylation status. This speculation is supported by a recent study (van Sluis et al. 2020).

### The number of R repeats is not related to rDNA methylation

Next, we tested the correlation between 45S rDNA methylation rate and the copy number of the R repeat that is associated with TTF1 (Grummt et al. 1986). Notably, Spearman correlation between the 45S rDNA methylation rate and R repeat copy number differed among individuals (21/32 individuals showed >0.05 false discovery rate). Even in samples with a clear correlation, the tendency was not consistent among individuals and both positive and negative correlations were observed (Supplemental Fig. S9, Supplemental Table S4). We speculate that the correlations observed in some samples were the effect of the correlation of contiguous copies; thus, they do not reflect a true correlation. Collectively, these observations suggest that the R repeat plays little role in the transcription of rRNA.

### There is no strong correlation between age, instability and methylation

Next, we sequenced two samples from young individuals (20s) and two from older individuals (70s) to investigate age-related changes in rDNA structure (Supplemental Table S1, row# 4-7). Between the Cas9-enriched young and old samples, no large-scale structural differences were observed. This is consistent with the previous finding that aging does not increase such differences (Caburet et al. 2005).

It has been proposed that the rate of methylation of 45S rDNAs is increased with age (Watada et al. 2020; Wang and Lemos 2019). We therefore tested this relationship using the two young samples and two older samples obtained by Oxford Nanopore sequencing. However, neither the rate of unmethylated copies nor the average methylation level of methylated copies was increased in the older samples (Supplemental Table S1). It should be noted that we did not analyze methylation status in the same individual over time and that the number of samples was not sufficient to draw conclusions, which may explain why our results were different from previous studies.

### Higher rDNA copy number, more methylated coding regions

We analyzed the relationship between rDNA copy number and methylation rate. To estimate the rDNA copy number per cell, we used the ratio of rDNA reads to the total reads in each sample. By this calculation, the rDNA copy number ranged from 250 to 700 copies per cell (Supplemental Table S5; Fig. 5F). These values showed good agreement with previously reported data derived by different methods, such as quantitative PCR and short-read high-throughput sequencing (Malinovskaya et al. 2018; Parks et al. 2018).

To analyze the relationship between rDNA copy number and methylation rate, we used only the HPGP data that were generated by the Human Pangenomics Reference Consortium (HPRC), which were all thought to be obtained around the same period and basecalled with Guppy 4.0.11 (see Methods). This was done to avoid artifacts caused by different library preparation and sequencing conditions. From the analysis of 23 HPGP samples, we found that the number of rDNA copies and the ratio of unmethylated copies per cell were negatively correlated (Pearson correlation, r = 0.749, *p* = 1.15 * 10^−5^). In other words, the number of unmethylated copies was roughly constant irrespective of rDNA copy number per cell (Pearson correlation, r = 0.07, *p* = 0.625) (Fig. 5F, right).

### rDNA instability is increased in progeroid syndrome

We also analyzed two cell lines derived from patients with progeroid syndrome: namely, Bloom syndrome patient B cells derived by EBV transformation, and Werner syndrome patient primary fibroblast cells. These syndromes are caused by mutations in the DNA repair machinery, which increases genome instability (Killen et al. 2009). A previous study using the Fiber FISH method suggested that the structure of rDNA is highly aberrant (~50% of total rDNA copies) in Werner syndrome patients (Caburet et al. 2005). Based on the Cas9-enriched Oxford Nanopore sequencing method, the rate of non-canonical (e.g., real palindrome) copies was 1.2% and 2.4% in Werner and Bloom patient cells, respectively. These values are much higher than those in the normal samples (~0.2%), but much lower than the previously reported value (~50%). In both progeroid syndrome samples, we found characteristic reads, including a duplication within the 45S rDNA that may create a non-canonical rRNA structure (Supplemental Fig. S10; Supplemental Table S1). Interestingly, these mutations were concentrated around the 7,000–14,000-nt region in the reference. Because they were relatively rare in the other samples (Supplemental Table S1, column 7), we speculate that they are genomic instability “hotspots”, where mutation is frequent in DNA repair compromised cells.

### Human pluripotent stem cells have a different methylation status

Next, we analyzed rDNA methylation status in human induced pluripotent stem cells (hiPSCs, 201B7) by Cas9-enriched sequencing. In ESCs and iPSCs, rDNA is thought to be globally unmethylated because of the high transcription activity (Gupta and Santoro 2020). Unexpectedly, however, around half of the 45S rDNAs were methylated and the IGS was heavily methylated in the iPSCs, similar to differentiated cells (Fig. 6A; Supplemental Fig. S11, see Discussion). We also tested iPSCs derived from Werner syndrome patient fibroblast cells (A0031) (Shimamoto et al. 2014). Similarly in these iPSCs, a substantial proportion of 45S rDNAs were methylated and the frequency was higher than in the original A0031 fibroblast cells (52% vs 43%, *p* ≈ 1.2 * 10^−6^, Fisher’s exact test, Fig. 6B), in contrast to the results of previous studies (Woolnough et al. 2016; Wang and Lemos 2019).

**Figure 6.**
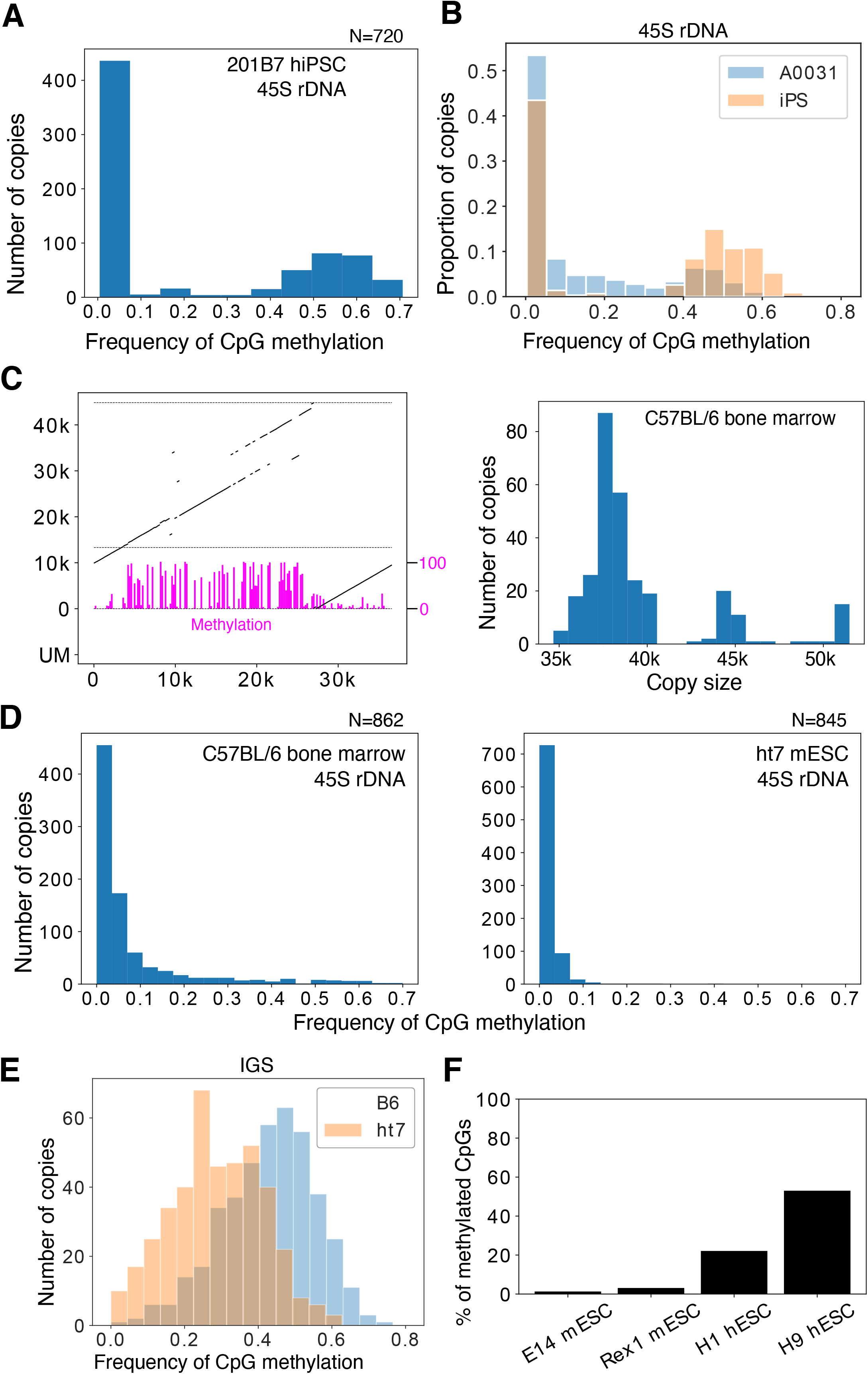
rDNA methylation of pluripotent stem cells. (A) The 45S rDNA methylation status of rDNA in hiPSCs does not differ from that in other differentiated samples: ~40% of the transcribed region is methylated. (B) Comparison of the 45S rDNA methylation status in Werner syndrome patient fibroblasts (A0031) and iPSCs derived from them. The *y* axis is the proportion of reads in each bin. The frequency of methylated copies is increased in iPSCs. (C) Representative rDNA from a Cas9-enriched Nanopore read of a mouse sample (left panel). Magenta bars represent CpG methylation. The estimated size distribution of mouse rDNA copies in C57BL/6 bone marrow cells is shown on the right. Note that the rDNA copy size of mouse is much smaller than previously reported. (D) 45S rDNA methylation levels in B6 bone marrow cells and ht7 mESCs. Methylation levels among copies are more continuous in mice and almost no methylation is seen in mESCs. (E) Comparison of IGS methylation levels in B6 bone marrow cells and ht7 mESCs. ht7 clearly shows lower a methylation level, even in the IGS region. (F) Proportion of CpGs methylated in the 45S rDNA of mESCs and hESCs determined by using publicly available short-read bisulfite whole genome sequencing data. While both samples of mESCs show a very low level of methylation, a substantial proportion of CpGs are methylated in hESCs.

In terms of rDNA stability, the frequency of aberrant structures was found to be significantly decreased in A0031-derived iPSCs (*p* ≈ 0.007, Fisher’s exact test, Supplemental Table S1). This is consistent with the finding that iPS induction suppresses chromosomal instability (Shimamoto et al. 2014), and we speculate that cells with stable genetic information are selected during the iPS induction process. If this is the case, the increased methylation of the transcribed region might be due to the selection process.

### rDNA structure and methylation in mouse

To test the generality of the Nanopore sequencer, we also analyzed two mouse-derived samples (bone marrow cells extracted from femur of an 8-week-old male C57BL/6J mouse and feeder-free ht7 embryonic stem cells (ESCs) derived from 129/Ola strain) (Niwa et al. 2000). Although mouse ribosomal DNA has been said to be around 45 kb in length (Grozdanov et al. 2003), we found that the average estimated size was much shorter than previous reported at 39.5 kb and 38 kb for C57BL/6J and ht7, respectively. Mouse rDNA also has variation in repeat number in the IGS, and this seems to determine the size distribution of rDNA (Fig. 6C; Supplemental Fig. S12).

In terms of methylation, 45S rDNAs of mESCs were almost completely free from methylation, which is comparatively different from hiPSCs (Fig. 6D right). In bone marrow cells, some of the 45S rDNAs were clearly methylated but, unlike in human, the distribution of CpG methylation among copies was continuous rather than bimodal, making it difficult to clearly define methylated copies (Fig. 6D left). The IGS was also methylated in mouse, but the frequency of methylation was much lower in mESCs than in hiPSCs (Fig. 6E). To examine whether the difference in methylation status between hiPSCs and mESCs was the result of the derivation method (i.e., ES vs iPS) or related to species differences (i.e., human vs mouse), we re-analyzed publicly available short-read-sequencer data, which showed that 45S rDNAs of mESCs are rarely methylated but a substantial proportion of CpGs in hESCs are methylated (Fig. 6F). Thus, the difference is likely to be ascribed to species variations. In fact, mESCs are known to be in a more undifferentiated state as compared with hESCs and show global hypomethylation (Nichols and Smith 2009; Nishino and Umezawa 2016).

## DISCUSSION

In this study, we analyzed the long rDNA array using data from the Oxford Nanopore sequencer. Our findings provide a new picture of the rDNA structure based on 39 samples with many reads that were directly extracted from human cells (~78,000 copies, more than 3 billion bases).

By using a new method that visualizes multiple copies of rDNA, we first characterized the rate of large-scale rDNA instability (inversion, deletion and other non-canonical structures) and found that such mutations are relatively rare, contrary to a previous report based on the Fiber FISH method (Caburet et al. 2005). We also showed that Oxford Nanopore sequencing occasionally produces artifactual palindromic reads that are considered to be difficult to distinguish from true palindromic reads (de Lannoy et al. 2018). Fortunately, we found such artificial palindromic reads have characteristic features, such as a poor quality score around the inversion and the position of the inversion site, and succeeded in recognizing them. As a result, we conclude that real palindromic structures are relatively rare. Lastly, we found that, although rDNA instability was not increased by age in our samples, it was increased in cells from patients with progeroid syndrome. In our study, ~40 samples were analyzed and only 0.2% (on average) of structures were non-canonical. This value is much lower (~1/30) than the value previously reported using Fiber FISH (Caburet et al. 2005). Thus, we conclude that the human rDNA is a relatively regular array.

We found that the R and Butterfly/Long repeat regions are variable in different copies, although they are similar in contiguous copies. Because the R and Butterfly/Long repeat regions have small repetitive sequences within the repeats, they may form a secondary DNA structure and inhibit the replication folk to induce instability. The R repeat contains many copies of the Sal box associated with TTF1 (Fig. 1A), which is known to arrest the replication fork both *in vivo* and *in vitro* (Santoro and Grummt 2005; Akamatsu and Kobayashi 2015). This sequence may work as a recombination hotspot, as observed in budding yeast (Supplemental Fig. S1; see below).

In contrast to variation, our study indicated that there is structural similarity between IGSs in contiguous copies. This suggests that gene conversion takes place frequently in the human rDNA, as in the budding yeast rDNA (Ganley and Kobayashi 2011). Previous studies also have suggested the possibility of IGS homogenization within the same chromosome (Gonzalez and Sylvester 2001). Our study strongly supports this view.

We also found that IGS are classified into two types among individuals (Fig. 3B; Supplemental Fig. S3). The mechanism behind this bimodal distribution of Butterfly/Long repeat copy number is a fascinating issue to be addressed in future studies. Another mystery is the presence of a rare type of IGS, which was observed in many samples. If the efficiency of gene conversion is high, such copies should be excluded. One explanation for the rare-type IGS is that these variations are often generated by chance and resolved over time by homogenization.

In the budding yeast, it is known that gene conversion occurs frequently and all copies essentially have the same sequence in a cell (Ganley and Kobayashi 2007). As a mechanism, Fob1/RFB-dependent rDNA recombination seems to be important (Ganley and Kobayashi 2011). As mentioned above, such an RFB site is also present in the R repeat in human rDNA. Therefore, a similar recombination repair system may also contribute to sequence homogenization in human cells (Akamatsu and Kobayashi 2015). Moreover, in progeroid syndrome patient cells, in which the activity of DNA repair is reduced, the number of non-canonical copies increased. Such rDNA instability has been also observed in the yeast WRN homologue mutant *sgs1* (Sinclair and Guarente 1997).

In terms of rDNA methylation, we found that there is an obvious difference between methylated and unmethylated 45S rDNAs (Fig. 5A,B). In unmethylated 45S rDNAs, the methylation rate was close to 0 such that the genes are likely to be transcribed (reviewed in Kass et al., 1997). The methylation status was also similar in contiguous copies. This may be because heterochromatin forms around these regions and affects rDNA silencing. In contrast to the “all-or-none” methylation pattern of 45S rDNAs, the non-coding IGS regions were always methylated, and the level was correlated with the methylation level of the contiguous 45S rDNAs (Fig. 5D). These observations suggest that the IGS is always in a similar heterochromatin structure and that 45S rDNA activation affects the region. We also have to note that, because all of the IGS regions in the rDNA are heavily methylated, the transcriptional rate of non-coding sequences in these regions may not be very high. Therefore, if this is the case, at least some non-coding transcripts are like to come from rDNA fragments that are scattered all over the human genome (Cherlin et al. 2020).

We found that unmethylated copies are negatively correlated with total rDNA copy number in a cell, suggesting that the number of unmethylated copies is roughly constant in different individuals, at least in the same tissue. This finding also supports our definition of unmethylated copies and our view that they are the actively transcribed ones. One possible mechanism behind the regulation of active copy number is that the transcription factor dosage is limited. Alternatively, the volume of the nucleolus fibrillar center may be kept at a constant level, which restricts the number of rDNA copies that can be held inside. Further analysis will be required to reveal the underlying mechanism.

In terms of mouse rDNA, we found that some features were similar to the human rDNA, such as repeat number variation in the IGS. One clear finding was that the unit length was shorter (~39 kb) than reported previously (~45 kb) (Grozdanov et al. 2003). The reason might be the difficulty in assembling sequences filled with repeats using relatively short reads.

Lastly, regarding the methylation in ES and iPS cells, the results differed between human and mouse. We expected that both cell types would be less methylated than their differentiated counterparts because the nucleolus of ES and iPS cells is known to be larger and rRNA transcription activity is high (Woolnough et al. 2016; Gupta and Santoro 2020; Wang and Lemos 2019). In the human iPSC and ESCs, however, the methylation status of the 45S rDNA was similar to that in differentiated cells. In contrast, most of the 45S rDNA copies in the mouse ESCs were unmethylated. This difference between mouse and human is thought to stem from the difference between their developmental stages. Mouse ESCs are always in a pre-X-chromosome-inactivated status and are globally hypomethylated; in human ESCs, by contrast, the X-chromosome is already inactivated in many cases and the other genomic regions are also highly methylated (Tomoda et al. 2012; Nishino and Umezawa 2016; Nichols and Smith 2009).

In summary, our results have revealed several new aspects of the most highly transcribed house-keeping gene, rDNA, in terms of its stability, structure, and methylation status.

## METHODS

### DNA extraction

DNA was extracted by modified Sambrook and Russell DNA extraction. In brief, 1 × 10^6^ cells were pelleted and 500 μL of TLB was added (10 mM Tris-Cl pH 8.0, 25 mM EDTA pH 8.0, 0.5% (w/v) SDS, 20 μg/ml RNase A). After mixing by inversion and incubating for 1 hour at 37 °C, 1 μL of 20 mg/ml Proteinase K (Roche) solution was added and the mixture was further incubated for 3 hours at 55 °C. Next, 500 μL of TE-saturated phenol was added and the solution was rotated until the water phase was clear. Phase lock gel (Dow Corning(R) High Vacuum Grease) was added, followed by centrifugation for 5 min, and then the upper phase was decanted into a 1.5-mL tube. Phenol/chloroform/isoamyl alcohol (25:24:1) was added and the above procedure was repeated. The resultant solution was placed in a 5-mL tube, and 200 μL of 5 M ammonium acetate and 1.5 mL of 100% ethanol were added with gentle rotation at RT until the solution was homogeneous. The DNA precipitate was collected by pipetting and placed in a 1.5 mL tube containing 70% ethanol. After centrifugation, 50 μL of TE was added to the pellet, which was left overnight at 4 °C. DNA concentration was measured with a Qubit assay kit (Invitrogen).

### Construction of the Cas9-enriched Oxford Nanopore library

The published protocol (Gilpatrick et al. 2020) was modified specifically for rDNA, which consists of hundreds of copies. First, Cas9 RNP was assembled as described previously. Then, 500 ng of DNA was dissolved in 9 μL of 1 × CutSmart buffer (New England Biolabs) and sheared by pipetting 30 times with a pipette set at 8 μL with a P2/P10 tip. One microliter of QuickCIP (New England Biolabs) was added and the solution was incubated at 37 °C for 10 min, followed by 80 °C for 2 min. Next, 0.5 μL of RNP, 0.3 μL of 10 mM dATP and 0.3 μL of Taq DNA polymerase were added to the solution before incubation at 37 °C for 15 min and 72 °C for 5 min. The ligation mix (5 μL of LNB, 3 μL of Quick ligase (New England Biolabs), 1.2 μL of MQ, 0.8 μL of AMX) was then added to the DNA solution in two stages, with tapping to mix between additions. After incubating for 10 min at room temperature, 2.7 μL of 5 M NaCl was added, followed by incubation for 5 min. After centrifuging at 15,000 rpm for 5 min, the supernatant was removed and 100 μL of 4.5% PEG 6000, 0.5 M NaCl, 5 mM Tris-HCl (pH 8.0) was added. After centrifuging again for 1 min, the supernatant was removed and the pellet was dissolved in 10 μL of EB. In our experience, centrifugation with salt rather than Ampure beads resulted in a higher library yield and a shorter centrifugation time was preferable. It was extremely rare to find reads containing more than two copies of rDNA with this method, indicating that the in vitro Cas9 efficiency is sufficient. The four gRNA target sequences were 5’-ATGAACCGAACGCCGGGTTAAGG, 5’-AGGACGGTGGCCATGGAAGTCGG, 5’-ACCTCCACCAGAGTTTCCTCTGG and 5’-TATCCTGAGGGAAACTTCGGAGG.

### Mice

Eight-week-old C57BL6/JJc1 mice were purchased from CLEA Japan, Inc (Tokyo, Japan). Bone marrow cells were extracted as described previously (Madaan et al. 2014). All experiments were approved by the Animal Experiment Ethics Committees of the University of Tokyo (Experiment No. 0210) and performed in accordance with the provided manual.

### Cell culture

EBV-transformed B cells were obtained from National Institutes of Biomedical Innovation, Health and Nutrition. 201B7 hiPSCs were obtained from Riken BioResource Research Center. EBV-transformed B cells were cultured as a floating culture in T25 flasks containing RPMI1640 supplemented with 10% FBS. hiPSCs were cultured on vitronectin-coated plates with AK02N medium. ht7 mESCs were cultured on 0.1% gelatin-coated plates with standard GMEM-based medium (10% FCS, 1xNEAA, 1 mM sodium pyruvate. 10^−4^ M 2-ME, 1000 U/mL mLIF). A0031 Werner syndrome patient cells and the iPSCs derived from them were cultured as described previously (Shimamoto et al. 2014).

### Screening of rDNA-derived Oxford Nanopore reads

To analyze the HPGP whole genome sequencing samples, we downloaded the fast5, fastq and sequencing summary files. First, based on the sequencing summary file, we excluded reads that did not pass the sequencing quality filtering. Next, each read was split and mapped to the rDNA reference file (KY962518.1) by using BWA MEM (v0.7.17) and the “ont2d” option. We only used reads that had more than 40,000 nt of continuous rDNA region at either end. Moreover, to remove reads derived from a microsatellite stretch similar to that included in the IGS of rDNA, we checked whether the reads contained at least 10% of split-reads that mapped to the coding region.

### Visualization of the rDNA-derived Oxford Nanopore reads

Each fastq read was split into smaller reads of 300-nt, and mapped to the rDNA reference sequence by using BWA MEM (v0.7.17) as described above. The split-reads were then visualized as lines based on their position in the original read. To visualize the Phred quality score, the score was binned in 200-nt bins and the mean score was plotted. Visualization of CpG methylation was done similarly by binning reads in 200-nt bins. For each bin, the frequency of methylation was calculated based on the “average” (see below) and the value was plotted as a bar.

### Finding non-canonical copies

First, the read was split and mapped to the rDNA reference sequence. If more than 10% of the mapped segment was in the opposite direction to the dominant direction, the read was classified as inverted and plotted. If the distance between each mapped read differed from the expected length by more than 500 nt, the reads were plotted as potential reads containing non-canonical copies. In case that both of two neighboring reads were within the R repeat or Butterfly/Long repeat region, we did not count them as aberrant copies owing to the natural variation in these regions. Each plotted read was then manually classified based on the visualization.

### Estimation of repeat length

Using BWA MEM software, 500-nt rDNA sections located at 10,000, 20,000 and 30,000 nt in the reference sequence were mapped to each read. Next, the distance between the mapped positions of 500-nt sections at 10,000 and 20,000 in each read was used to estimate the R repeat length and the distance between the mapped positions of 20,000 and 30,000 was used to estimate the Butterfly/Long repeat length. Because genomic mutations in rDNA are rare, most of the variations obtained by this method should be due to repeat length variation in the repeat regions.

### Methylation analysis

For the reads that were thought to contain rDNA, fast5 files were extracted by using ont_fast5_api and basecalled by using Guppy Basecaller v4.2.2 with dna_r9.4.1_450bps_modbases_dam-dcm-cpg_hac_prom configuration. For threshold-based methylation analysis, we used 0.8 as the threshold for posterior probability. In the HPGP database, there are two types of Nanopore data, which were generated by NHGRI-USCS and HPRC. For the analysis comparing transcribed-region methylation frequency and rDNA copy number, we used only data generated by HPRC because they were submitted to the database over a short period of time and thus were likely to be less affected by differences in experimental conditions. The number of samples available was sufficient for the analysis (23 samples).

### Whole genome bisulfite sequencing analysis

Fastq files were first cleaned up with Trim Galore! (v0.6.6) to remove adapters (Martin 2011). The frequency of CpG methylation in the rDNA coding region was then estimated by Bismark software (v0.22.3) (Krueger and Andrews 2011) and Bowtie2 (v2.3.5) using the reference genome that contained only the rDNA coding region sequence. We used the following data: m14 mESC (SRR610046, SRA), Rex1 mESC (SRR5099302, SRA), H1 hESC (ENCFF311PSV, ENCODE project) and H9 hESC (ENCFF384QMG, ENCODE project). Files are available through SRA (https://www.ncbi.nlm.nih.gov/sra/) and ENCODE (https://www.encodeproject.org/files/), respectively.

## DATA ACCESS

All of the raw Cas9-enriched data generated in this study have been uploaded to Mendeley Data (https://dx.doi.org/10.17632/h48hj39bpm.1, https://dx.doi.org/10.17632/2wdg439sx4.1, https://dx.doi.org/10.17632/m84pty74mk.1).

## COMPETING INTEREST STATEMENT

The authors declare that they have no conflict of interest.

## ACKNOWLEDGEMENTS

We thank the members of Kobayashi lab for their useful discussion. This work was supported by AMED-CREST under grant number JP20gm1110010 to T.K.

## Notes

### Competing Interest Statement

The authors have declared no competing interest.

### Summary of Updates

Supplemental files were added.

## REFERENCES

Agrawal S, Ganley ARD. 2018. The conservation landscape of the human ribosomal RNA gene repeats. PLoS One 13: 1–31.

Akamatsu Y, Kobayashi T. 2015. The Human RNA Polymerase I Transcription Terminator Complex Acts as a Replication Fork Barrier That Coordinates the Progress of Replication with rRNA Transcription Activity. Mol Cell Biol 35: 1871–1881.

Burkhalter MD, Sogo JM. 2004. rDNA enhancer affects replication initiation and mitotic recombination: Fob1 mediates nucleolytic processing independently of replication. Mol Cell 15: 409–421.

Caburet S, Conti C, Schurra C, Lebofsky R, Edelstein SJ, Bensimon A. 2005. Human ribosomal RNA gene arrays display a broad range of palindromic structures. Genome Res 15: 1079–1085.

Carrero D, Soria-Valles C, López-Otín C. 2016. Hallmarks of progeroid syndromes: Lessons from mice and reprogrammed cells. DMM Dis Model Mech 9: 719–735.

Cherlin T, Magee R, Jing Y, Pliatsika V, Loher P, Rigoutsos I. 2020. Ribosomal RNA fragmentation into short RNAs (rRFs) is modulated in a sex- and population of origin-specific manner. BMC Biol 18: 1–19.

de Lannoy C, Ridder D De, Risse J. 2018. The long reads ahead : de novo genome assembly using the MinION. F1000Research 6: 1–26.

Defossez P-A, Prusty R, Kaeberlein M, Lin S-J, Ferrigno P, Silver PA, Keil RL, Guarente L. 1999. Elimination of Replication Block Protein Fob1 Extends the Life Span of Yeast Mother Cells. Mol Cell 3: 447–455. doi:https://doi.org/10.1016/S1097-2765(00)80472-4.

Gangloff S, Zou H, Rothstein R. 1996. Gene conversion plays the major role in controlling the stability of large tandem repeats in yeast. EMBO J 15: 1715–1725.

Ganley ARD, Kobayashi T. 2007. Highly efficient concerted evolution in the ribosomal DNA repeats: total rDNA repeat variation revealed by whole-genome shotgun sequence data. Genome Res 17: 184–191.

Ganley ARD, Kobayashi T. 2011. Monitoring the rate and dynamics of concerted evolution in the ribosomal DNA repeats of saccharomyces cerevisiae using experimental evolution. Mol Biol Evol 28: 2883–2891.

Ganley ARD, Kobayashi T. 2014. Ribosomal DNA and cellular senescence: new evidence supporting the connection between rDNA and aging. FEMS Yeast Res 14: 49–59.

Gilpatrick T, Lee I, Graham JE, Raimondeau E, Bowen R, Heron A, Downs B, Sukumar S, Sedlazeck FJ, Timp W. 2020. Targeted nanopore sequencing with Cas9-guided adapter ligation. Nat Biotechnol 38: 433–438.

Gonzalez IL, Sylvester JE. 2001. Human rDNA: Evolutionary patterns within the genes and tandem arrays derived from multiple chromosomes. Genomics 73: 255–263.

Grozdanov P, Georgiev O, Karagyozov L. 2003. Complete sequence of the 45-kb mouse ribosomal DNA repeat: Analysis of the intergenic spacer. Genomics 82: 637–643.

Grummt I, Rosenbauer H, Niedermeyer I, Maier U, Öhrlein A. 1986. A repeated 18 bp sequence motif in the mouse rDNA spacer mediates binding of a nuclear factor and transcription termination. Cell 45: 837–846.

Gupta S, Santoro R. 2020. Regulation and Roles of the Nucleolus in Embryonic Stem Cells: From Ribosome Biogenesis to Genome Organization. Stem Cell Reports 15: 1206–1219.

Kaeberlein M, McVey M, Guarente L. 1999. The SIR2/3/4 complex and SIR2 alone promote longevity in Saccharomyces cerevisiae by two different mechanisms. Genes Dev 13: 2570–2580.

Kass SU, Landsberger N, Wolffe AP. 1997. DNA methylation directs a time-dependent repression of transcription initiation. Curr Biol 7: 157–165.

Killen MW, Stults DM, Adachi N, Hanakahi L, Pierce AJ. 2009. Loss of Bloom syndrome protein destabilizes human gene cluster architecture. Hum Mol Genet 18: 3417–3428.

Kim JH, Dilthey AT, Nagaraja R, Lee HS, Koren S, Dudekula D, Wood WH, Piao Y, Ogurtsov AY, Utani K, et al. 2018. Variation in human chromosome 21 ribosomal RNA genes characterized by TAR cloning and long-read sequencing. Nucleic Acids Res 46: 6712–6725.

Kobayashi T. 2008. A new role of the rDNA and nucleolus in the nucleus - RDNA instability maintains genome integrity. BioEssays 30: 267–272.

Kobayashi T. 2011. Regulation of ribosomal RNA gene copy number and its role in modulating genome integrity and evolutionary adaptability in yeast. Cell Mol Life Sci 68: 1395–1403.

Kobayashi T. 2014. Ribosomal RNA gene repeats, their stability and cellular senescence. Proc Japan Acad Ser B Phys Biol Sci 90: 119–129.

Kobayashi T. 2003. The Replication Fork Barrier Site Forms a Unique Structure with Fob1p and Inhibits the Replication Fork. Mol Cell Biol 23: 9178–9188.

Kobayashi T, Ganley ARD. 2005. Recombination regulation by transcription-induced cohesin dissociation in rDNA repeats. Science (80-) 309: 1581–1584.

Kobayashi T, Heck DJ, Nomura M, Horiuchi T. 1998a. Expansion and contraction of ribosomal DNA repeats in Saccharomyces cerevisiae: requirement of replication fork blocking (Fob1) protein and the role of RNA polymerase I. Genes Dev 12: 3821–3830.

Kobayashi T, Heck DJ, Nomura M, Horiuchi T. 1998b. Expansion and contraction of ribosomal DNA repeats in Saccharomyces cerevisiae: Requirement of replication fork blocking (Fob1) protein and the role of RNA polymerase I. Genes Dev 12: 3821–3830.

Kobayashi T, Horiuchi T, Tongaonkar P, Vu L, Nomura M. 2004. SIR2 regulates recombination between different rDNA repeats, but not recombination within individual rRNA genes in yeast. Cell 117: 441–453.

Krueger F, Andrews SR. 2011. Bismark: A flexible aligner and methylation caller for Bisulfite-Seq applications. Bioinformatics 27: 1571–1572.

Li H. 2013. Aligning sequence reads, clone sequences and assembly contigs with BWA-MEM. arXiv 00: 1–3.

Madaan A, Verma R, Singh AT, Jain SK, Jaggi M. 2014. A stepwise procedure for isolation of murine bone marrow and generation of dendritic cells. J Biol Methods 1: 1.

Malinovskaya EM, Ershova ES, Golimbet VE, Porokhovnik LN, Lyapunova NA, Kutsev SI, Veiko NN, Kostyuk S V. 2018. Copy number of human ribosomal genes with aging: Unchanged mean, but narrowed range and decreased variance in elderly group. Front Genet 9: 306.

Martin M. 2011. Cutadapt removes adapter sequences from high-throughput sequencing reads. EMBnet.journal 17: 10.

Miga KH, Koren S, Rhie A, Vollger MR, Gershman A, Bzikadze A, Brooks S, Howe E, Porubsky D, Logsdon GA, et al. 2020. Telomere-to-telomere assembly of a complete human X chromosome. Nature 585: 79–84.

Nichols J, Smith A. 2009. Naive and Primed Pluripotent States. Cell Stem Cell 4: 487–492.

Nishino K, Umezawa A. 2016. DNA methylation dynamics in human induced pluripotent stem cells. Hum Cell 29: 97–100.

Niwa H, Miyazaki JI, Smith AG. 2000. Quantitative expression of Oct-3/4 defines differentiation, dedifferentiation or self-renewal of ES cells. Nat Genet 24: 372–376.

Parks MM, Kurylo CM, Dass RA, Bojmar L, Lyden D, Vincent CT, Blanchard SC. 2018. Variant ribosomal RNA alleles are conserved and exhibit tissue-specific expression. Sci Adv 4: eaao0665.

Petes TD. 1979. Yeast ribosomal DNA genes are located on chromosome XII. Proc Natl Acad Sci U S A 76: 410–4.

Roussel P, André C, Comai L, Hernandez-Verdun D. 1996. The rDNA transcription machinery is assembled during mitosis in active NORs and absent in inactive NORs. J Cell Biol 133: 235–246.

Saka K, Ide S, Ganley ARD, Kobayashi T. 2013. Cellular senescence in yeast is regulated by rDNA noncoding transcription. Curr Biol 23: 1794–1798.

Santoro R, Grummt I. 2005. Epigenetic Mechanism of rRNA Gene Silencing: Temporal Order of NoRC-Mediated Histone Modification, Chromatin Remodeling, and DNA Methylation. Mol Cell Biol 25: 2539–2546.

Schawalder J, Paric E, Neff NF. 2003. Telomere and ribosomal DNA repeats are chromosomal targets of the bloom syndrome DNA helicase. BMC Cell Biol 4.

Shafin K, Pesout T, Lorig-Roach R, Haukness M, Olsen HE, Bosworth C, Armstrong J, Tigyi K, Maurer N, Koren S, et al. 2020. Nanopore sequencing and the Shasta toolkit enable efficient de novo assembly of eleven human genomes. Nat Biotechnol 38: 1044–1053.

Shimamoto A, Kagawa H, Zensho K, Sera Y, Kazuki Y, Osaki M, Oshimura M, Ishigaki Y, Hamasaki K, Kodama Y, et al. 2014. Reprogramming suppresses premature senescence phenotypes of Werner syndrome cells and maintains chromosomal stability over long-term culture. PLoS One 9: 1–13.

Sinclair DA, Guarente L. 1997. Extrachromosomal rDNA Circles— A Cause of Aging in Yeast. Cell 91: 1033–1042.

Takeuchi Y, Horiuchi T, Kobayashi T. 2003. Transcription-dependent recombination and the role of fork collision in yeast rDNA. Genes Dev 17: 1497–1506.

Tomoda K, Takahashi K, Leung K, Okada A, Narita M, Yamada NA, Eilertson KE, Tsang P, Baba S, White MP, et al. 2012. Derivation conditions impact X-inactivation status in female human induced pluripotent stem cells. Cell Stem Cell 11: 91–9.

van Sluis M, van Vuuren C, Mangan H, McStay B. 2020. NORs on human acrocentric chromosome p-arms are active by default and can associate with nucleoli independently of rDNA. Proc Natl Acad Sci U S A 117: 10368–10377.

Wang M, Lemos B. 2019. Ribosomal DNA harbors an evolutionarily conserved clock of biological aging. Genome Res 29: 325–333.

Watada E, Li S, Hori Y, Fujiki K, Shirahige K, Inada T, Kobayashi T. 2020. Age-Dependent Ribosomal DNA Variations in Mice. Mol Cell Biol 40.

Weitao T, Budd M, Hoopes LLM, Campbell JL. 2003. Dna2 helicase/nuclease causes replicative fork stalling and double-strand breaks in the ribosomal DNA of Saccharomyces cerevisiae. J Biol Chem 278: 22513–22522.

Woolnough JL, Atwood BL, Liu Z, Zhao R, Giles KE. 2016. The regulation of rRNA gene transcription during directed differentiation of human embryonic stem cells. PLoS One 11: 1–18.

