## Supplemental Figures S1-S12 for "The human ribosomal RNA gene is composed of highly homogenized tandem clusters"

**Supplemental Tables (Excel files)**

**Table S1. Number of non-canonical rDNA copies in each sample**

**Table S2. Methylation in contiguous rDNA copies**

**Table S3. Estimation of rDNA copy number**

**Table S4. 45S rDNA methylation and R repeat number**

**Table S5. Estimation of rDNA copy number per cell**

**Supplemental Figures**

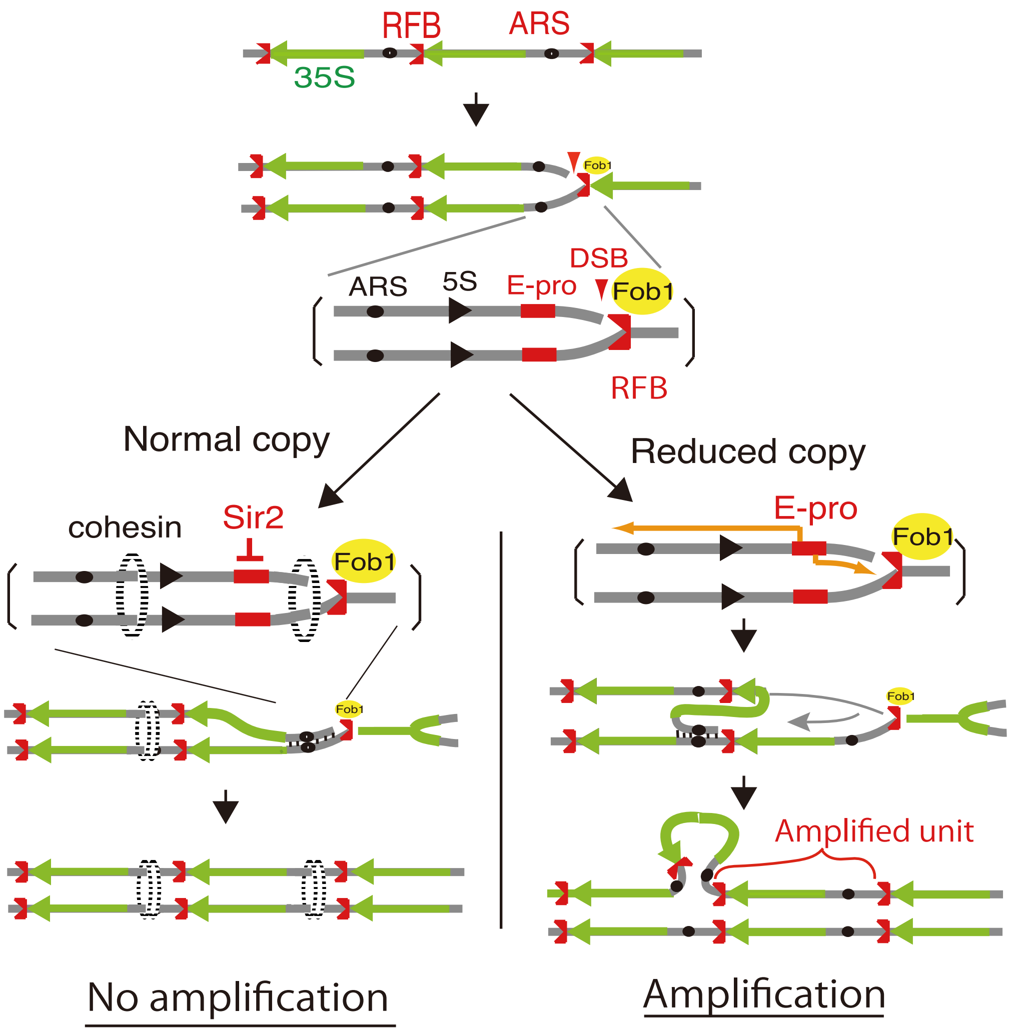


**Supplemental Figure S1.**

rDNA recombination for amplification in budding yeast.

Three copies of rDNA are shown. Green arrows represent the 35S rDNA and black arrowheads represent the 5S rDNA. The replication fork barrier (RFB), replication origin (ARS) and non-coding promoter (E-pro) are shown. Fob1 inhibits the replication fork and induces a DNA double strand break to trigger amplification of rDNA. Sir2 represses E-pro and ensures cohesin association, which inhibits unequal sister chromatid recombination for amplification.

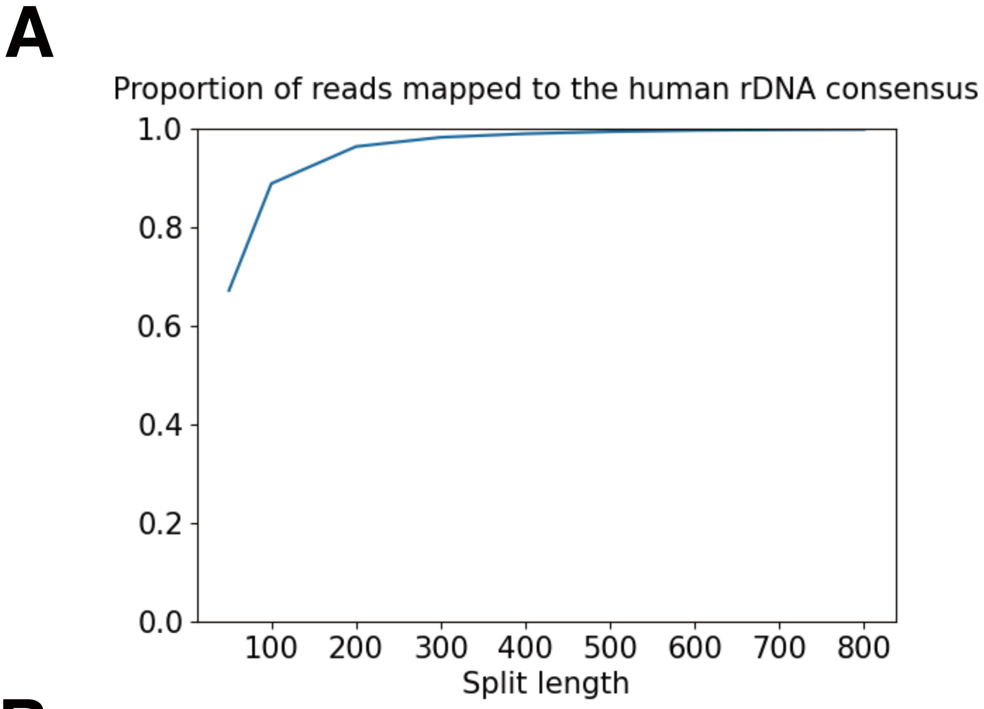


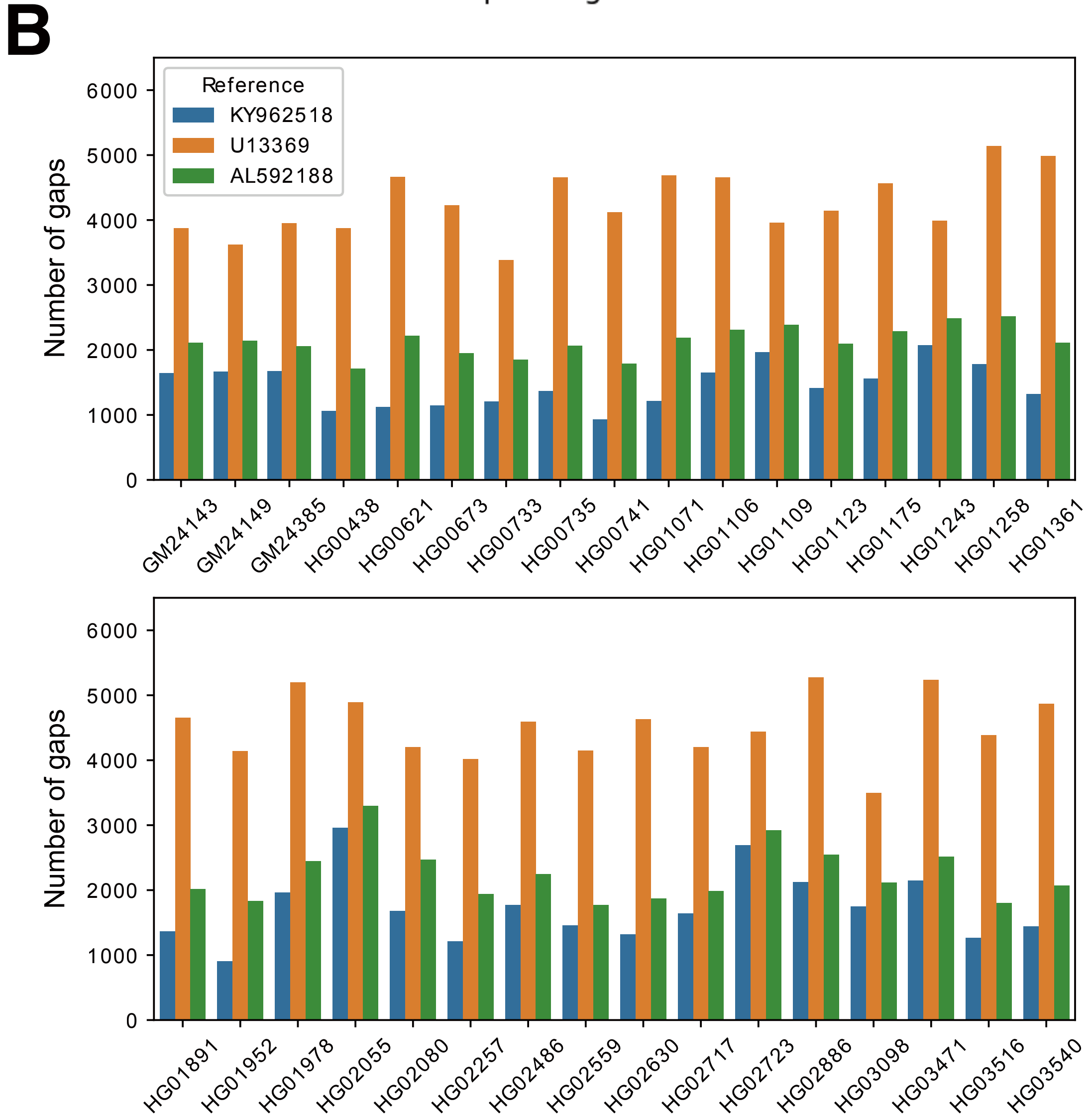


**Supplemental Figure S2**

(A) Proportion of Nanopore reads that mapped to the human rDNA consensus sequence by split length. Longer split length resulted in higher mapping frequency.

(B) Split reads mapped to three different reference rDNA sequences. The distances between the mapped positions of successive split reads were calculated. If the distances differed from the expected distance by more than 100 nt, they were counted as “gaps” . We counted the number of these gaps to determine which reference was the most similar to actual rDNA sequences.

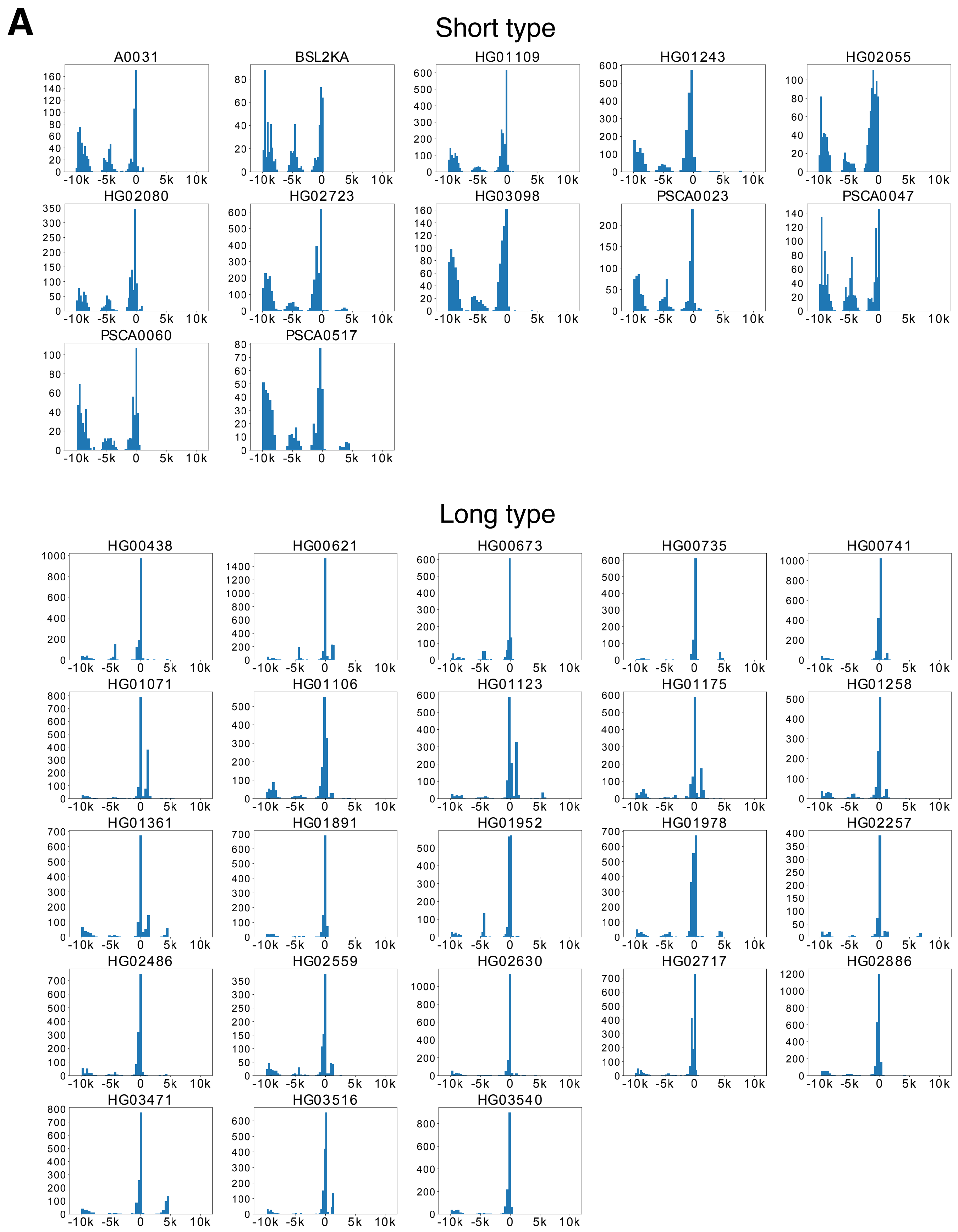


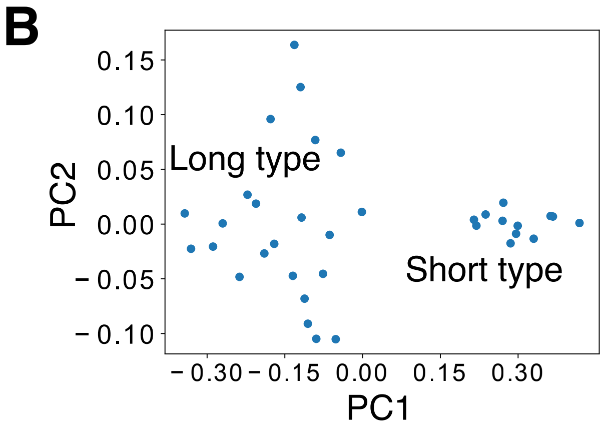


**Supplemental Figure S3**

(A) Length distributions of the Butterfly/Long repeat region in various samples. The distribution could be roughly classified into two types (short and long) based on average length with each type sharing a similar distribution.

(B) Principal component analysis (PCA) of the Butterfly/Long repeat distribution. Each distribution was vectorized by binning by repeat length and by calculating the proportion of copies falling into each bin. The distributions could clearly be distinguished simply by the first component.

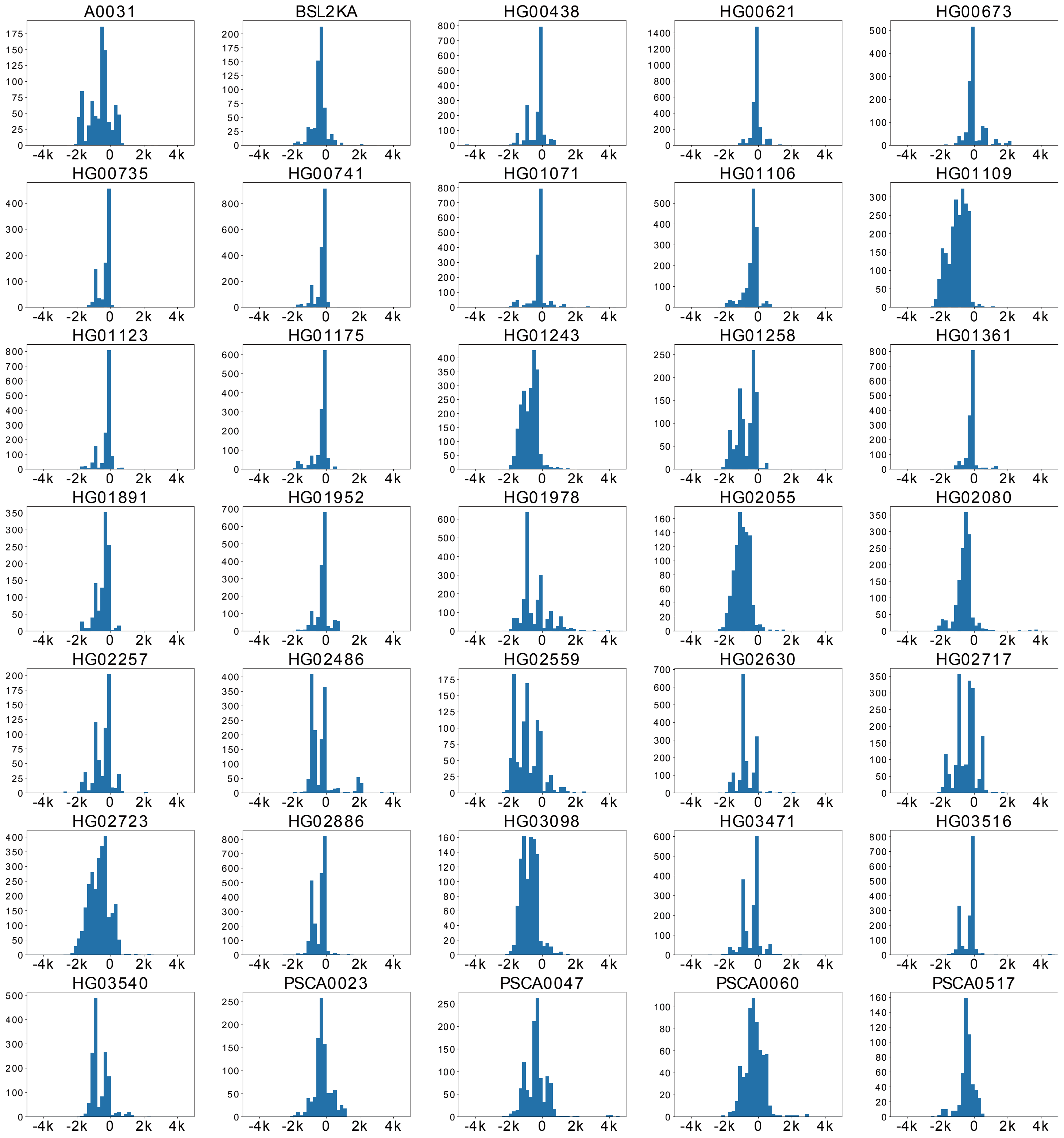


**Supplemental Figure S4**

Length distribution of the R repeat region in various samples. Peaks corresponding to different numbers of repeats are evident in some samples, while the distribution is more continuous in others. Overall, there is a considerable variation among samples.

### Butterfly/long repeat

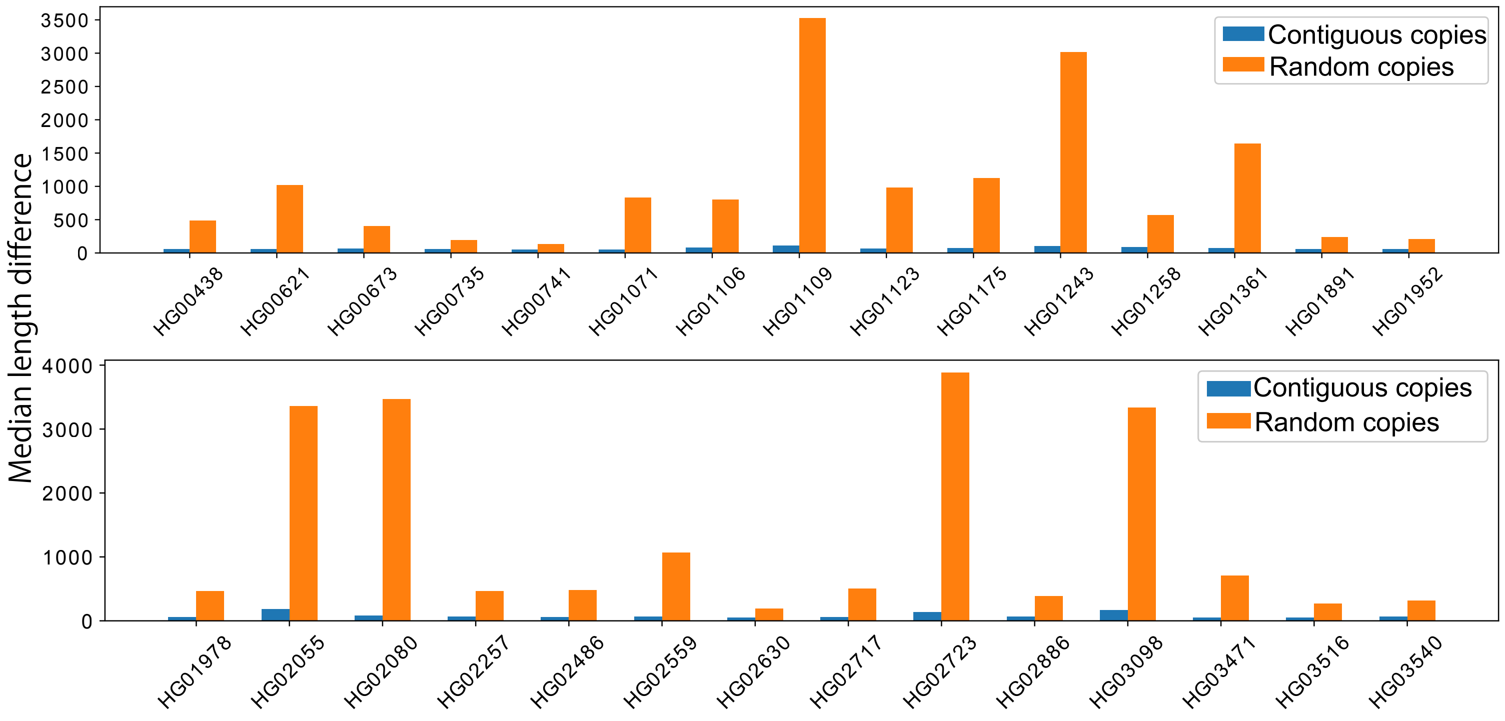


### R repeat

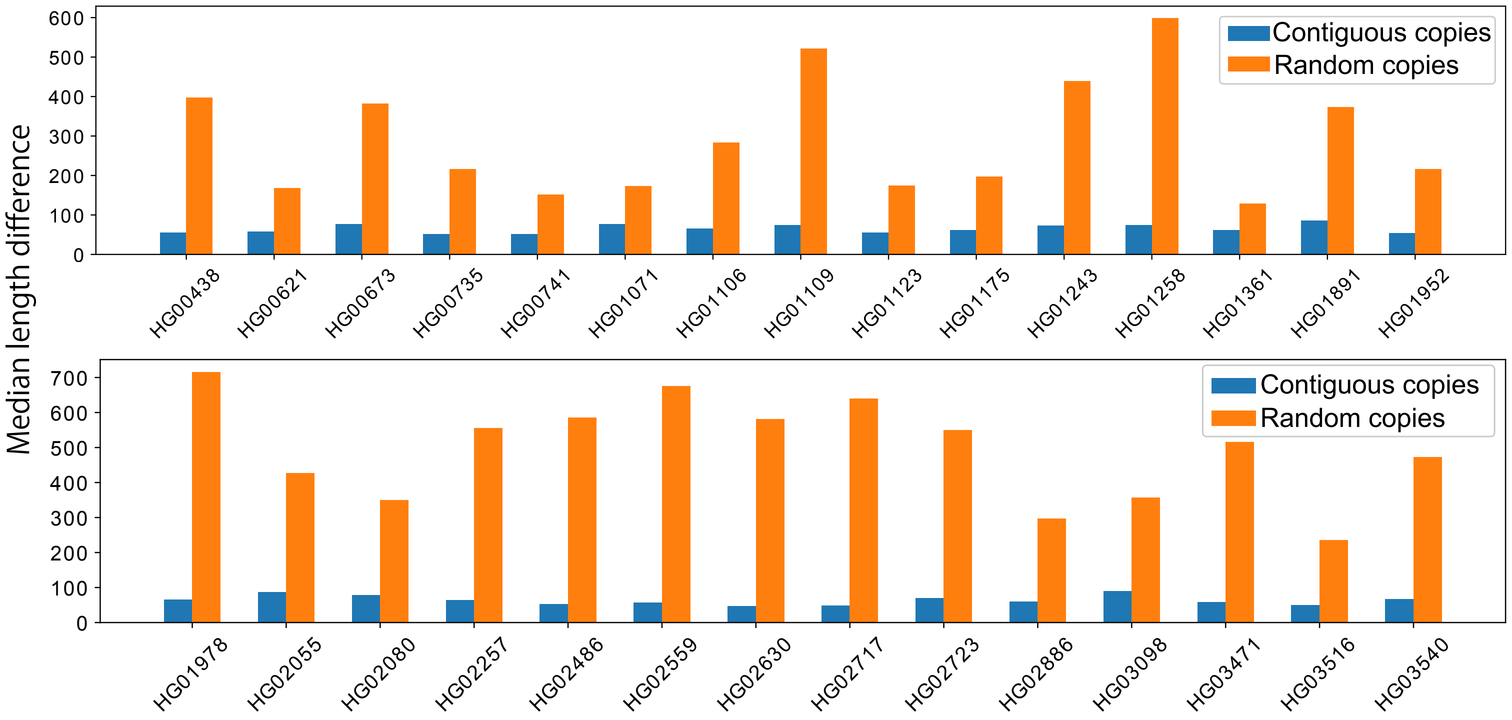


**Supplemental Figure S5**

Comparison of the median difference in length of the Butterfly/long repeat and R repeat between contiguous and random copies. Contiguous copies are clearly similar to each other.

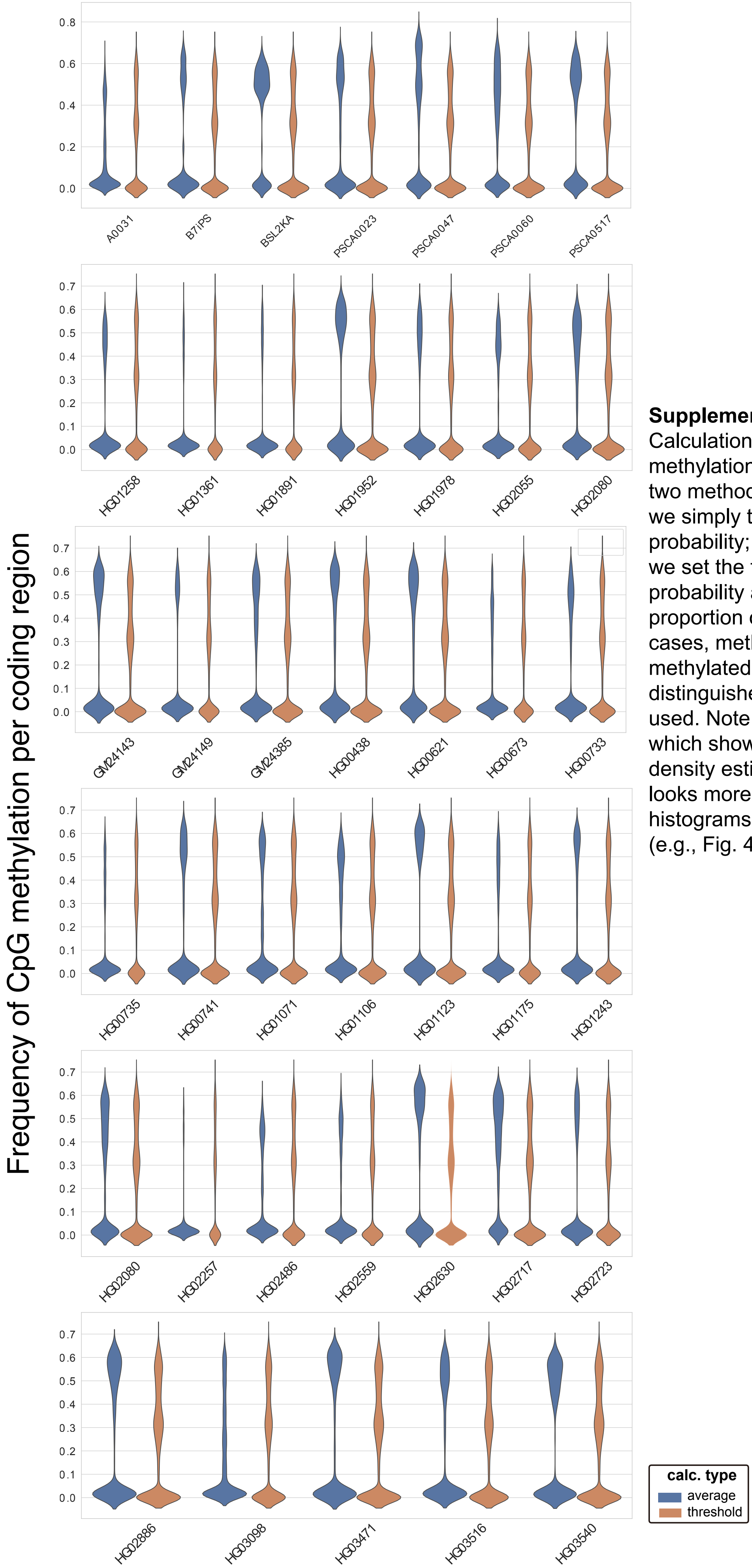


**Supplemental Figure S6**
Calculation of the frequency of CpG methylation in the rDNA coding region by two methods. In the "average" method (blue), we simply took the mean of the posterior probability; in the "threshold" method (orange), we set the threshold of the posterior probability at 0.8 and calculated the proportion of methylated CpGs. In most cases, methylated (>0.2) and less methylated (~0.0) copies could easily be distinguished no matter which method was used. Note that here we used a violin plot, which shows data summarized by kernel density estimation; therefore, the distribution looks more continuous than the actual histograms shown in the main manuscript (e.g., Fig. 4B).

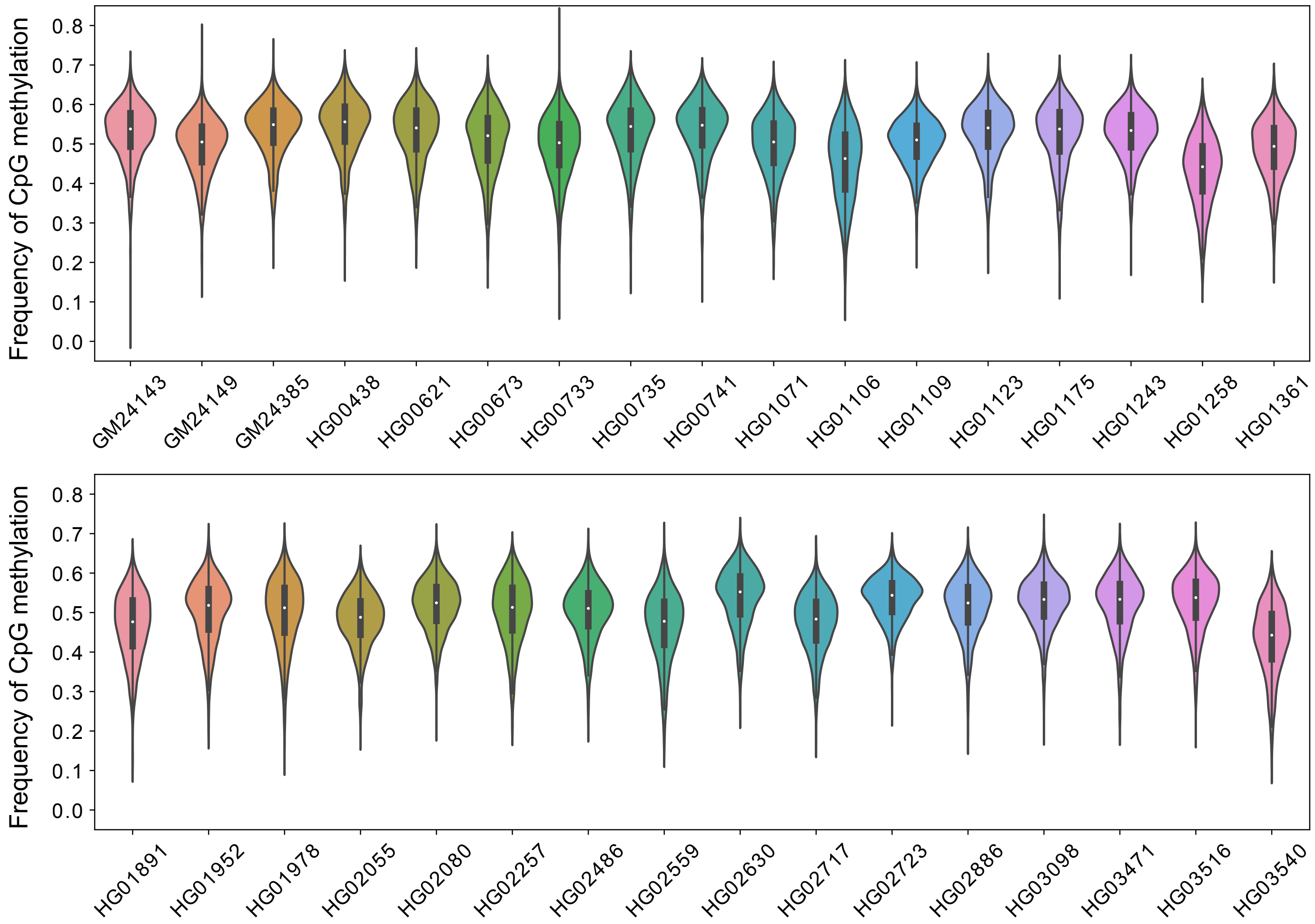


**Supplemental Figure S7**

Methylation levels of the IGS region in different individuals. The frequency of methylation in all copies was above that of the 45S unmethylated cutoff, 0.05.

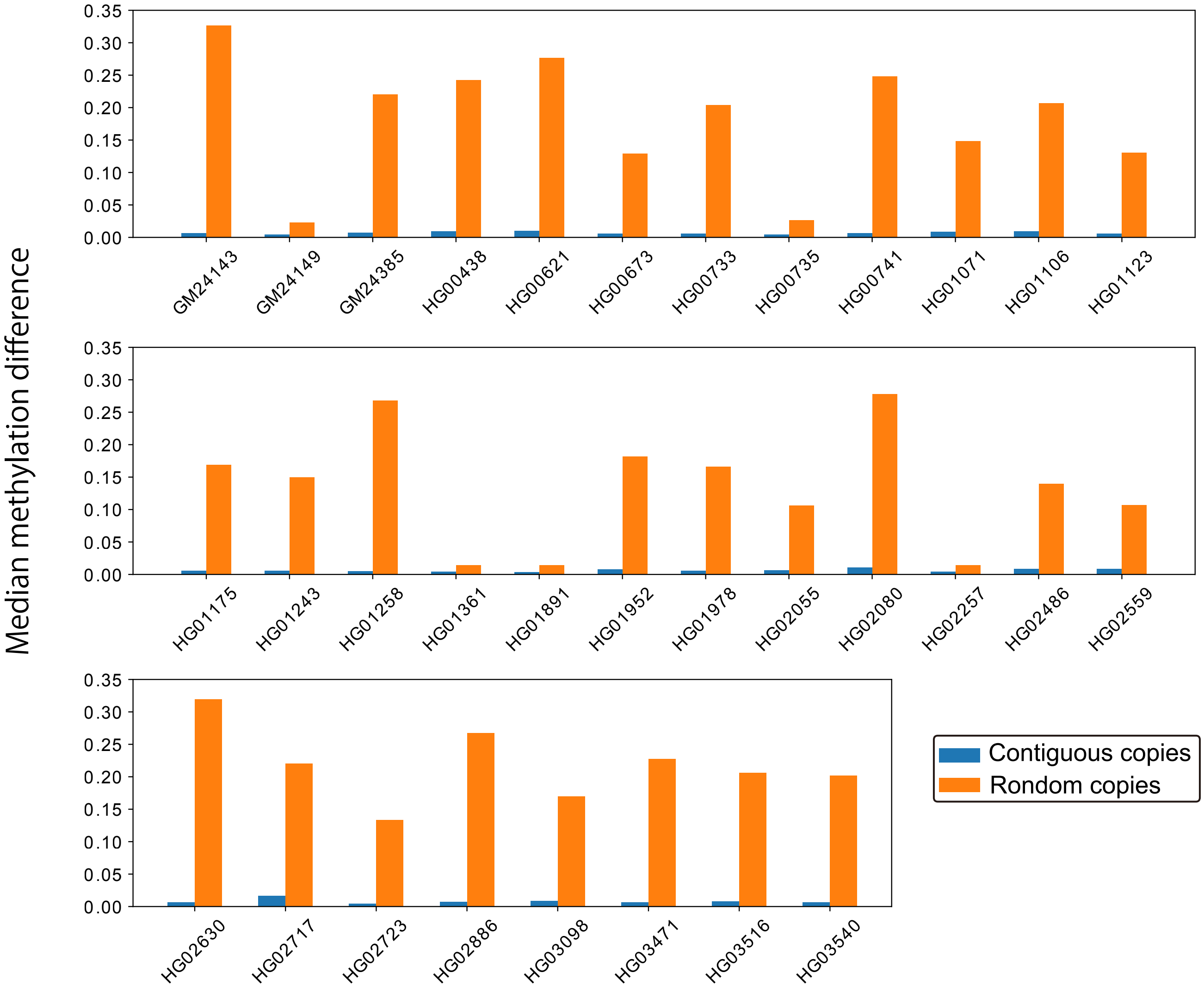


**Supplemental Figure S8**

Comparison of the median difference in CpG methylation level in 45S rDNA between contiguous and random copies. Contiguous copies share a similar methylation level regardless of sample.

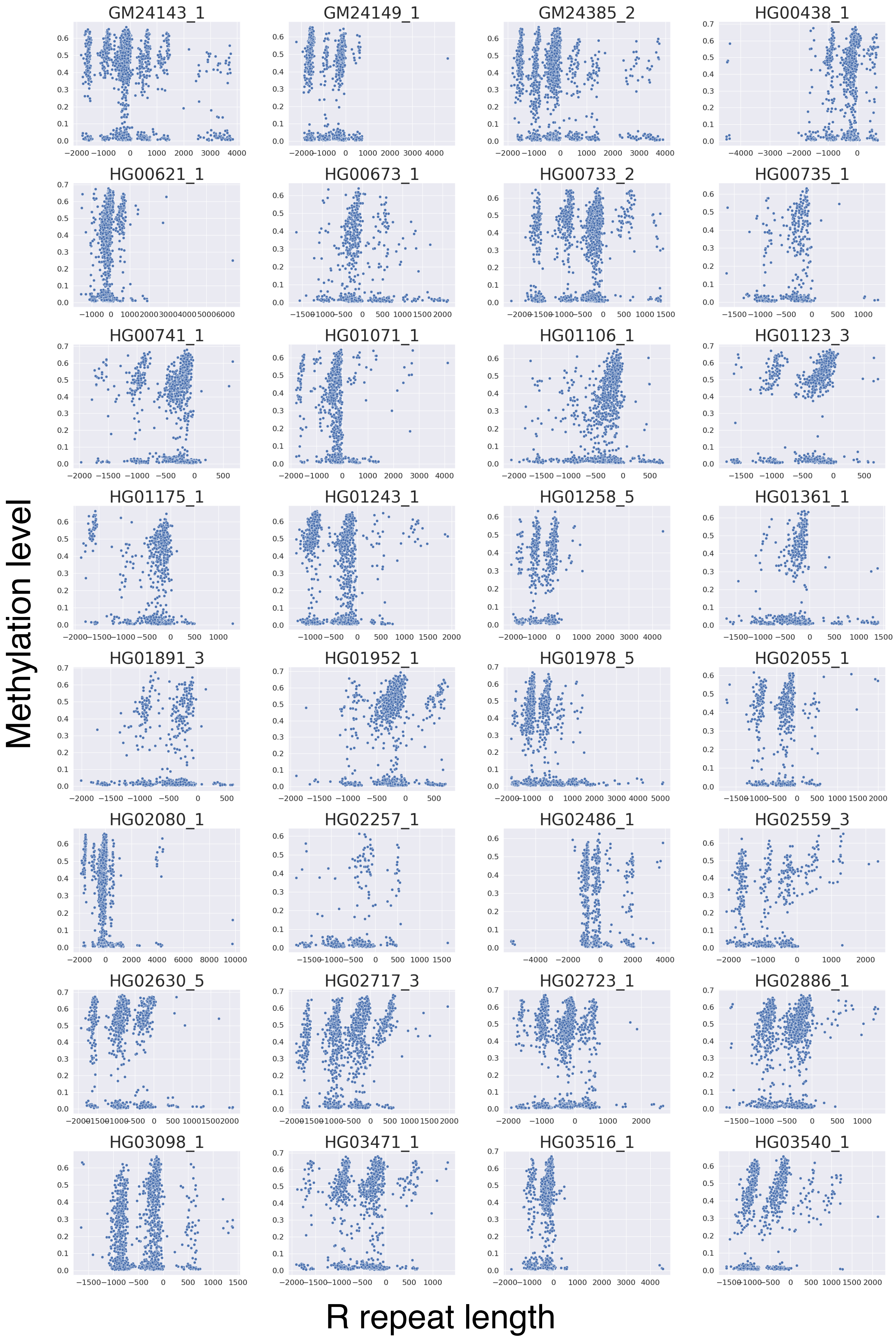


**Supplemental Figure S9**

Relationship between the methylation level in 45S rDNA and the length of contiguous R repeats. There may be some correlations, but they differ among individuals; that is, there is no consensus relationship. Therefore, any apparent correlations may be byproducts of correlations between contiguous copies.

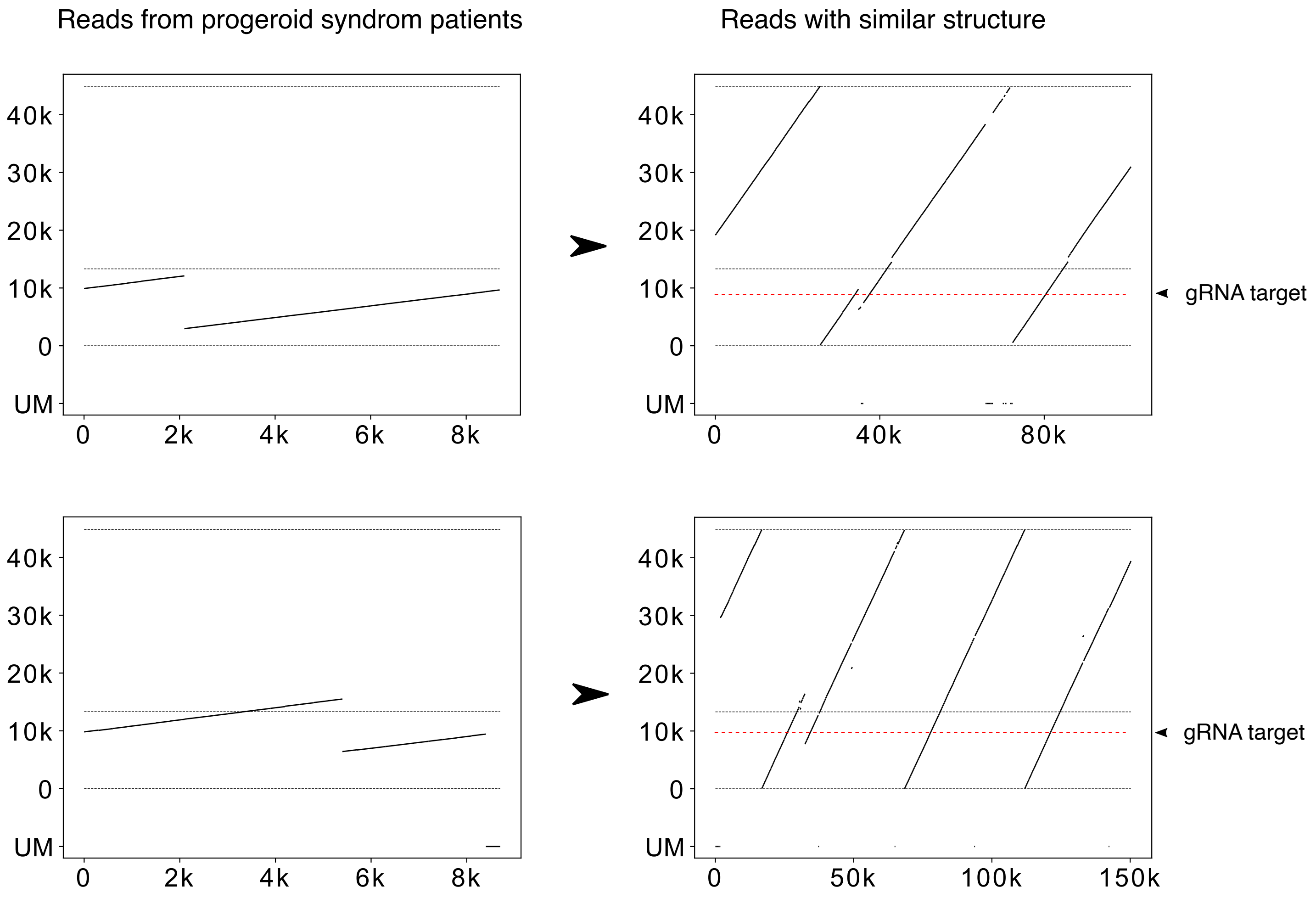


**Supplemental Figure S10**

(Left panels) Reads from Cas9-enriched samples. (Right panels) Examples of WGS reads with mutations that have recombinational hotspots similar to those seen in Cas9-enriched progeroid syndrome samples. UM: unmapped

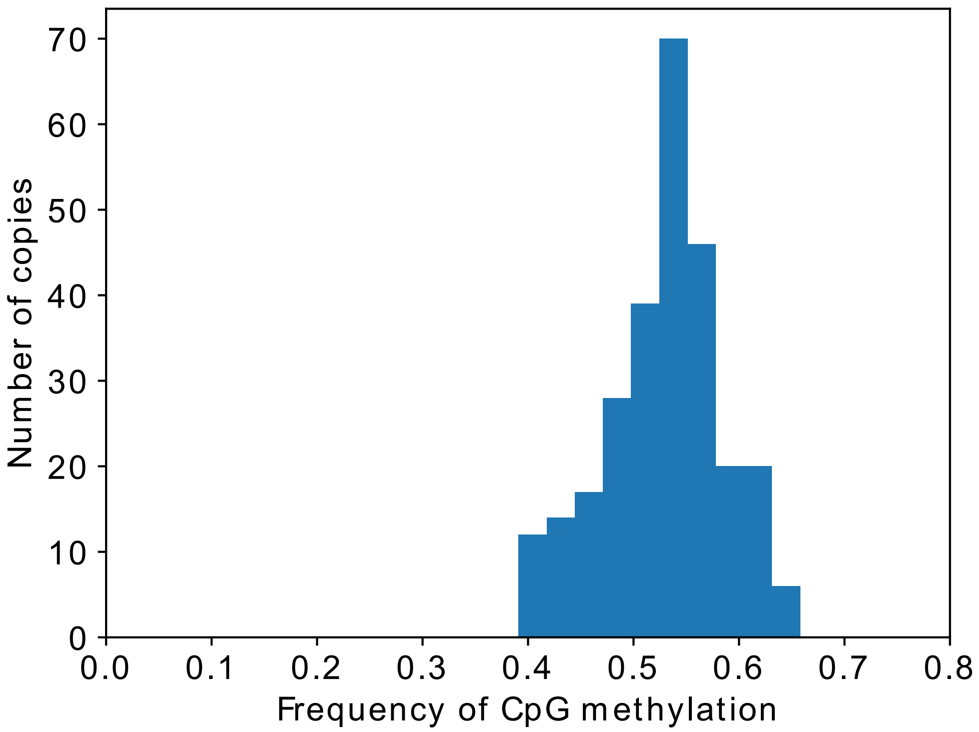


**Supplemental Figure S11**

Methylation level of the IGS in 201B7 hiPSCs.

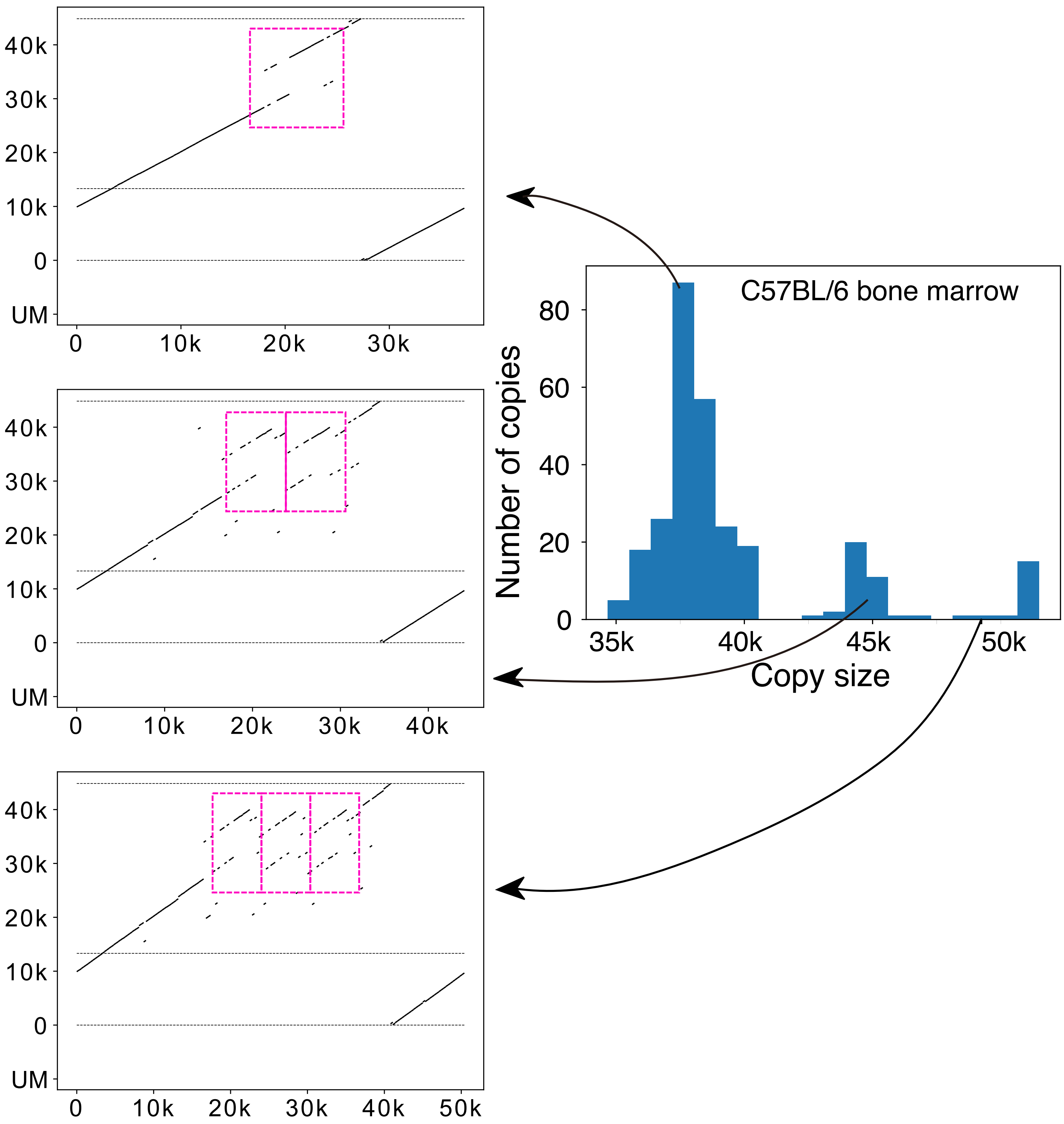


**Supplemental Figure S12**
Distribution of rDNA copy size in mouse C57BL/6J bone marrow cells and three representative visualizations of the rDNA in each peak. Repeat units in the IGS are indicated by dashed squares.
